# A MAP kinase cascade broadly regulates development and virulence of *Sclerotinia sclerotiorum* and can be targeted by HIGS for disease control

**DOI:** 10.1101/2023.03.01.530680

**Authors:** Lei Tian, Josh Li, Yan Xu, Yilan Qiu, Xin Li

**Author notes:** These authors contributed equally to this work (Lei Tian, Josh Li).

## Abstract

*Sclerotinia sclerotiorum* causes white mold or stem rot in a broad range of economically important plants, bringing significant yield losses worldwide. Host-induced gene silencing (HIGS) has been showing promising effects in controlling many fungal pathogens, including *S. sclerotiorum*. However, molecular genetic understanding of signaling pathways involved in its development and pathogenicity is needed to provide effective host-induced gene silencing (HIGS) targets for disease control. Here, by employing a forward genetic screen, we characterized an evolutionarily conserved mitogen-activated protein kinase (MAPK) cascade in *S. sclerotiorum*, consisting of SsSte50-SsSte11-SsSte7-Smk1, controlling mycelial growth, sclerotia development, compound appressoria formation, virulence, and hyphal fusion. Moreover, disruption of the putative downstream transcription factor SsSte12 led to normal sclerotia but aberrant appressoria formation and host penetration defects, suggestive of diverged regulation downstream of the MAPK cascade. Most importantly, targeting of *SsSte50* using host-expressed HIGS double stranded RNA resulted in largely reduced virulence of *S. sclerotiorum* on *Nicotiana benthamiana* leaves. Therefore, this MAPK signaling cascade is generally needed for its growth, development, and pathogenesis, and is an ideal HIGS target for mitigating economic damages caused by *S. sclerotiorum* infection.

## Introduction

*Sclerotinia sclerotiorum* (Lib.) de Bary is a notorious fungal pathogen with an extremely wide host range. This ascomycete can infect more than 600 plant species, especially dicotyledons (Boland & Hall, 1994; Liang & Rollins, 2018). As one of the most devastating soilborne phytopathogens, it can infect important crops including canola, sunflower, soybean, tomato, lettuce, and onion, causing severe yield losses worldwide (Boland & Hall, 1994; Bolton et al., 2006). Like other fungal pathogens, *S. sclerotiorum* hyphae can differentiate into penetration compound appressoria structure to perforate the plant epidermal layer (Tariq & Jeffries, 1984). During infection, the key virulence factor oxalic acid (OA) is produced, which induces cell wall degradation and host cell death (Liang et al., 2015). Upon nutrient depletion, *S. sclerotiorum* can form sclerotia, resting structures which can resist freezing temperatures, abiotic stresses, and many fungicides (Bolton et al., 2006; Xia et al., 2020). In the field, sclerotia can overwinter and germinate under favorable conditions carpogenically to release airborne ascospores or myceliogenically to directly infect neighboring plants (Erental et al., 2008). Despite its economic importance, the detailed molecular mechanisms for *S. sclerotiorum* development and pathogenicity are not well understood.

Eukaryotes rely on signal transduction pathways to respond to environmental changes and coordinate complex cellular functions. The highly conserved mitogen-activated protein kinase (MAPK) cascades are broadly used for regulating gene expression, cell proliferation, and differentiation (Meloche & Pouyssegur, 2007; Cargnello & Roux, 2011). In budding yeast *Saccharomyces cerevisiae*, several MAPK cascades function in parallel to regulate growth, cell wall integrity, osmoregulation, and reproduction (Hamel et al., 2012). Among them, the pheromone response pathway is responsible for recognition of extracellular mating factors, and the signal transduction occurs via sequential phosphorylation mediated through the activation of the following components: the MAPKKK Ste11 with its adaptor protein Ste50, the MAPKK Ste7, and the final MAPK Kss1/Fus3. Activated Kss1/Fus3 subsequently phosphorylates the transcription factor Ste12, which promotes the expression of genes involved in the formation of copulatory structures (Chen & Thorner, 2007; Hamel et al., 2012). Moreover, the MAPKKK Ste11 and the adaptor protein Ste50 are also involved in high osmolarity response through activating the MAPKK Pbs2 and the MAPK Hog1 sequentially in yeast (Posas & Saito, 1997; Zarrinpar et al., 2004; Hamel et al., 2012).

In phytopathogenic fungi, parallel MAPK signaling pathways are highly conserved and used in more complex biological processes such as pathogenesis and appressoria development. For example, in rice blast *Magnaporthe oryzae*, the homologous Ste50-Ste11-Ste7-Kss1/Fus3 cascade also exists. Deletion of orthologs of Ste7 and Ste11 completely abolished appressoria formation required for host penetration (Zhao et al., 2005). Likewise, the deletion *M. oryzae* Mst50 (Ste50 homolog) yielded appressoria with decreased turgor pressure, attenuating epithelial penetration (Cai et al., 2022). In Sclerotiniaceae *Botrytis cinerea*, the BMP1 (Botrytis MAP kinase required for Pathogenesis) MAPK cascade shares homologous kinases as the *S. cerevisiae* pheromone response pathway (Sharma & Kapoor, 2017). Disruption of components of the cascade results in retarded growth, loss of pathogenicity and reduced conidia formation (Zheng et al., 2000; Doehlemann et al., 2006; Schamber et al., 2010). These cases demonstrate the conservation of the MAPK cascades in fungi, and their crucial roles in regulating growth, development, and pathogenicity.

Although parallel *S. cerevisiae* MAPK cascades in *S. sclerotiorum* were identified through homology comparison with yeast, few were characterized molecularly (Hegedus et al., 2016). The disruption of the cell wall integrity MAPK pathway components, including Pkc1, Bck1, Mkk1, and Smk3, not only led to decreased cell wall integrity, but also decreased sclerotia formation and virulence (Bashi et al., 2016; Cong et al., 2022). Likewise, deletion of the MAPKKK gene *Ssos4*, a component of the high osmolarity MAPK pathway associated with mitigating hyperosmotic stressors, led to reduced sclerotia formation and virulence (Li et al., 2021). SsSte12, the homolog of *S. cerevisiae* pheromone responsive transcription factor Ste12 activated by MAPK Kss1/Fus3, was studied through RNA interference (RNAi)-mediated gene knockdown in *S. sclerotiorum*. *Ssste12* RNAi mutants with reduced gene expression exhibited slower growth, smaller sclerotia and fewer appressoria (Xu et al., 2018). However, the exact function of the parallel pheromone response MAPK cascade in *S. sclerotiorum* remains largely unclear.

In this study, we report the identification of *SsSte50* and *SsSte11* from our forward genetic screen searching for mutants with defective sclerotia (Xu et al., 2022). Next generation sequencing (NGS) data analysis revealed several candidate mutations. Gene knockout and transgene complementation confirmed the causal mutations in *SsSte50* and *SsSte11*, which encode an MAPK adaptor and a MAPKKK respectively in the same MAPK cascade. Furthermore, we showed that deletion of the MAPKK gene *SsSte7* or MAPK gene *Smk1* phenocopies *Ssste50* and *Ssste11* mutants with no sclerotia and compound appressoria formation, and are non-pathogenic, supporting orthologous MAPK cascade to yeast. However, knocking out the putative downstream transcription factor SsSte12 leads to normal sclerotia but aberrant compound appressoria formation and host penetration defects. Finally, we observed largely enhanced resistance of *Nicotiana benthamiana* plants against *S. sclerotiorum* upon expression of host-induced gene silencing (HIGS) dsRNA targeting *SsSte50*. Therefore, components from the MAPK signaling cascade can be targeted for mitigating economic damages caused by *S. sclerotiorum* infection.

## Results

### Three ultraviolet (UV)-induced *S. sclerotiorum* mutants exhibit similar defects in vegetative growth, sclerotia formation and virulence

To identify genes that are involved in sclerotia development in *S. sclerotiorum*, we performed a UV mutagenesis-based forward genetic screen searching for mutants with no sclerotia formation (Xu et al., 2022). When compared with the wild-type (WT) *S. sclerotiorum* strain 1980, three mutants shared almost identical defects including thickened mycelia and slower mycelial growth were identified: Z-2, MT-4 and R7 (Figure 1A-B).

**Figure 1.**
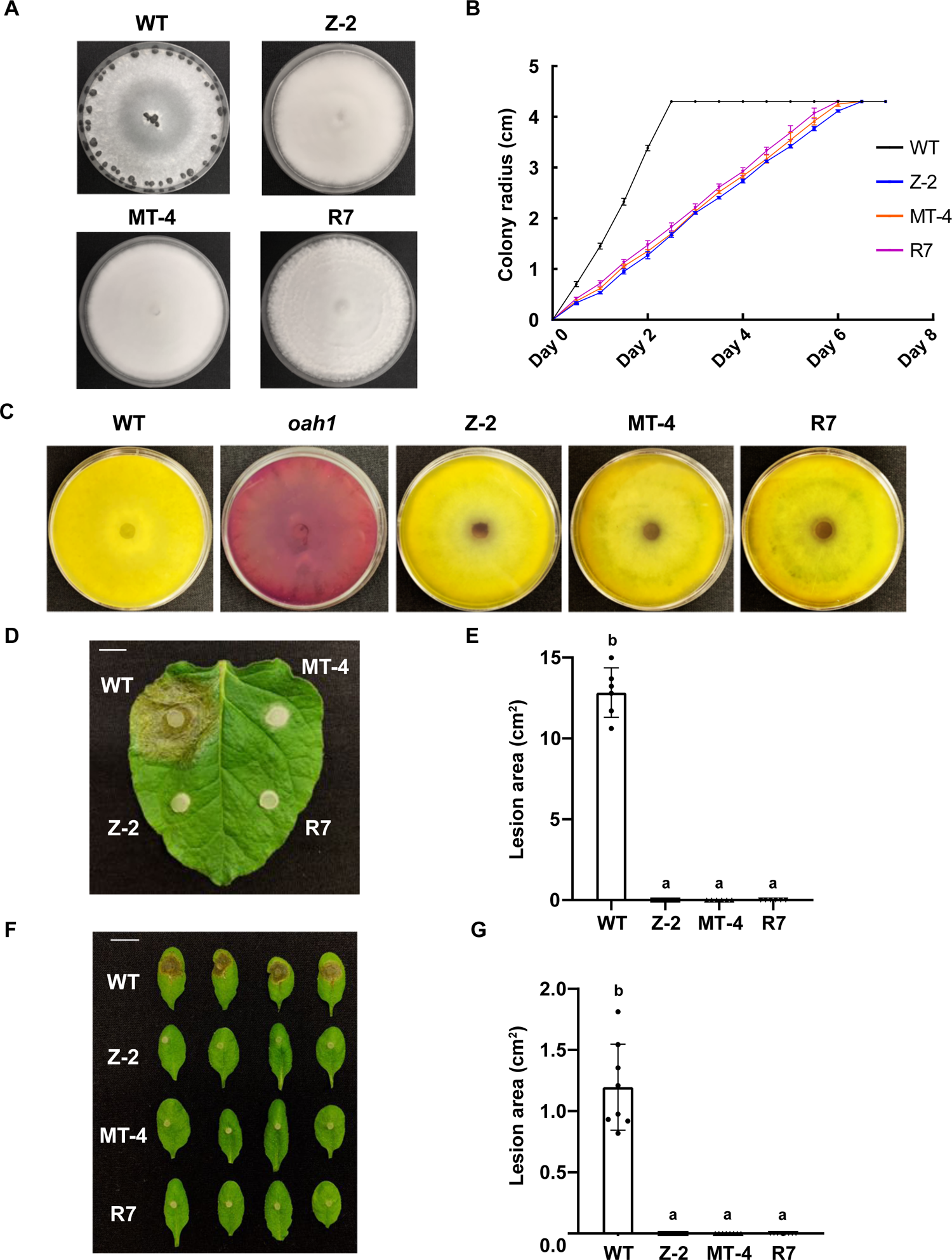
Phenotypic characterization of three *S. sclerotiorum* UV mutants Z-2, MT-4, and R7. A. Colony morphology of wild-type (WT) strain 1980 and the UV mutants Z-2, MT-4, and R7 on potato dextrose agar (PDA) plates. The pictures were taken at 14 days post inoculation (dpi). B. Mycelial growth of WT, Z-2, MT-4, and R7 on PDA plates. The colony radius was measured every 12 hours until the mycelia reach the plate edge. Two independent experiments were carried out with similar results. C. Plate images of WT, *oah1*, Z-2, MT-4, and R7 grown on PDA media supplemented with 50 mg/L bromophenol blue (BB). Oxalic acid (OA) production is indicated by the change in PDA-BB medium from violet to yellow. The pictures were taken at 4 dpi. Three independent experiments were carried out with similar results. The OA deficient mutant *oah1* serves as a positive control. D. Pathogenicity test of WT, Z-2, MT-4, and R7 on detached *N. benthamiana* leaves. The picture was taken at 36 hours post inoculation (hpi). E. Quantification of the lesion areas caused by the indicated *S. sclerotiorum* strains at 36 hpi. Statistical significance is indicated by different letters (*p* < 0.01). Error bars represent means ± SD (*n* = 6). Three independent experiments were carried out with similar results. Bar = 1 cm. F. Pathogenicity test of WT, Z-2, MT-4, and R7 on detached *A. thaliana* Col-0 leaves. The picture was taken at 36 hpi. G. Quantification of the lesion areas caused by the indicated *S. sclerotiorum* strains at 36 hpi. Statistical significance is indicated by different letters (*p* < 0.01). Error bars represent means ± SD (*n* = 8). Three independent experiments were carried out with similar results. Bar = 1 cm.

To test for virulence alterations, we first assayed oxalic acid (OA) production using potato dextrose agar (PDA) plates with the pH indicator bromophenol blue (pH > 4.6 - violet; pH < 3.0 - yellow), as OA is a key virulence factor in *S. sclerotiorum* (Liang *et al*., 2015). WT, Z-2, MT-4, and R7 plates changed color from violet to yellow while the control plates inoculated with the OA biosynthesis mutant *oah1* (*<u>O</u>xaloacetate <u>A</u>cetyl<u>H</u>ydrolase <u>1</u>*) (Liang *et al*., 2015) remained violet, indicating that the OA production was not affected in all three mutants (Figure 1C). When tested on detached WT *N. benthamiana* and *Arabidopsis thaliana* Col-0 ecotype leaves, all three mutants failed to cause any necrotic lesions at 36 hours post-inoculation (hpi) (Figure 1D-G). Even at 4 days post-inoculation when the whole leaves were macerated by WT *S. sclerotiorum*, no lesion was detected with the mutants, indicating that the loss of virulence is not only due to growth defects (Supplemental Figure S1). Collectively, as Z-2, MT-4, and R7 exhibited highly similar defects in all assays, we suspected that the proteins encoded by their causal genes may participate in the same biological pathway, contributing to multiple processes in Sclerotinia biology including growth, sclerotia formation and virulence.

### Different mutations in the MAPK adaptor protein-encoding gene *SsSte50* are responsible for Z-2, MT-4, and R7 mutant phenotypes

To identify the causal mutations for the Z-2, MT-4 and R7 phenotypes, genomic DNA from each of the mutants was extracted and sequenced via whole genome next-generation sequencing (NGS). The NGS data analysis was completed with an established pipeline (Xu *et al*., 2022). Interestingly, these mutants possess different frameshift mutations in *sscle_07g058440*, suggesting that it is likely the gene whose mutations lead to Z-2, MT-4, and R7 phenotypes (Supplemental Figure S2). Protein structural analysis using AlphaFold revealed that Z-2, MT-4, and R7 carry frameshift mutations in the SAM (Sterile Alpha Motif) domain, the downstream alpha helix, and the Ras-associating domain, respectively (Figure 2A). R7 contains a more downstream mutation compared to Z-2 and MT-4, explaining some phenotypic leakiness seen in R7. For instance, sclerotia initiation hyphal aggregates were observed in R7, though they could not develop into mature sclerotia (Fig. 1A).

**Figure 2.**
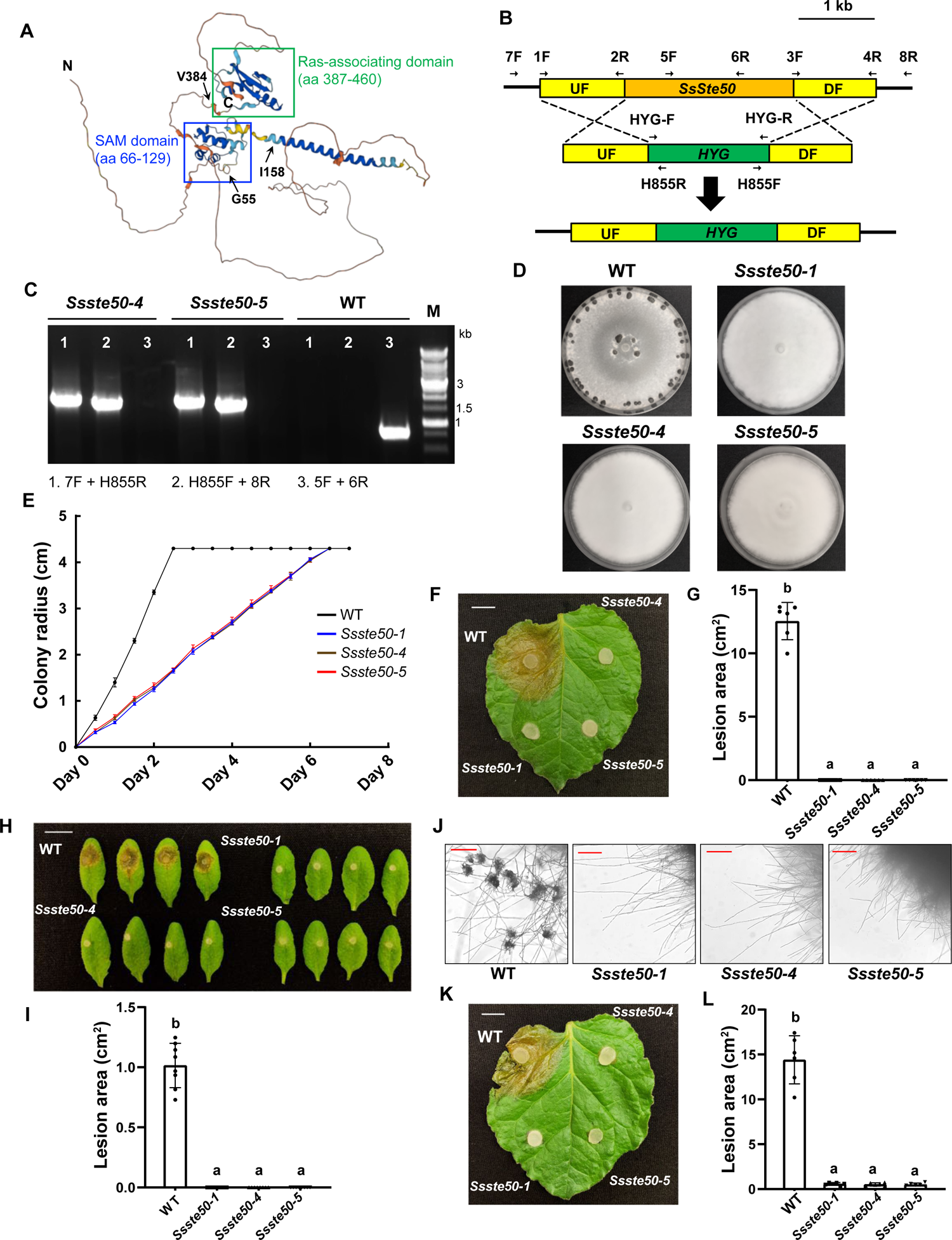
Knocking out *SsSte50* phenocopies Z-2 (renamed as *SsSte50-1*). A. Protein structure of SsSte50 as predicted by AlphaFold. Two main domains, SAM (Sterile Alpha Motif) domain and Ras-associating domain are highlighted in blue and green boxes respectively with amino acid (aa) positions labeled below. Frameshift (fs) mutations in Z-2 (G55fs), MT-4 (I158fs), and R7 (V384fs) are indicated with arrows. N: amino (N)-terminus; C: carboxyl (C)-terminus. B. Target gene knock-out (KO) strategy through homologous recombination. The *SsSte50* and hygromycin-resistance gene *HYG* are shown as orange and green rectangles respectively. The primers specified with arrows were used to amplify the flanking regions or to genotype the KO transformants. UF: upstream fragment; DF: downstream fragment. C. PCR verification of *SsSte50* gene KO. The genomic DNA from WT *S. sclerotiorum* and two KO mutants *Ssste50-4*, *Ssste50-5* were used as PCR templates. Primer pairs 1 and 2 were used to test the *HYG* gene insertion, and primer pair 3 was used to test the deletion of *SsSte50*. Lane M is the DNA size ladder. D. Colony morphology of WT, *SsSte50-1*, *Ssste50-4*, and *Ssste50-5* on PDA plates. The pictures were taken at 14 dpi. E. Mycelial growth of the indicated genotypes on PDA plates. The radius was measured every 12 hours until the fungus colonizes the whole plate. Two independent experiments were carried out with similar results. F. Pathogenicity test of the indicated genotypes on detached *N. benthamiana* leaves. The picture in F was taken at 36 hpi. G. Quantification of the lesion areas caused by the indicated *S. sclerotiorum* strains at 36 hpi. Statistical significance is indicated by different letters (*p* < 0.01). Error bars represent means ± SD (*n* = 6). Three independent experiments were carried out with similar results. Bar = 1 cm. H. Pathogenicity test of the indicated genotypes on detached *A. thaliana* Col-0 leaves. The picture in H was taken at 36 hpi. I. Quantification of the lesion areas caused by the indicated *S. sclerotiorum* strains at 36 hpi. Statistical significance is indicated by different letters (*p* < 0.01). Error bars represent means ± SD (*n* = 8). Three independent experiments were carried out with similar results. Bar = 1 cm. J. Compound appressoria formation by the indicated mutants. The mutants were transferred from PDA plates onto glass slides for compound appressoria observation. The pictures were taken at 2 dpi. The Bar = 200 μm. K. Pathogenicity test of the indicated genotypes on detached *N. benthamiana* leaves with pre-inoculation wounding. The picture in K was taken at 30 hpi. L. Quantification of the lesion areas caused by the indicated *S. sclerotiorum* strains at 30 hpi. Statistical significance is indicated by different letters (*p* < 0.01). Error bars represent means ± SD (*n* = 6). Three independent experiments were carried out with similar results. Bar = 1 cm.

Interestingly, *sscle_07g058440* is a single-copy gene with high homology to budding yeast *S. cerevisiae Ste50* (Supplemental Figure S3), which encodes an adaptor protein in the pheromone response MAPK pathway (Chen & Thorner, 2007; Hegedus et al., 2016). The involvement of Ste50 orthologs in *M. oryzae* and *B. cinerea* pathogenesis further suggests that the *sscle_07g058440* mutations are likely the causal ones for Z-2, MT-4 and R7 (Schamber et al., 2010; Cai et al., 2022). We thus named *sscle_07g058440* as *SsSte50*. Z-2, MT-4 and R7 were renamed as *SsSte50-1*, *SsSte50-2*, *SsSte50-3*.

To confirm that the *SsSte50* mutations are responsible for the mutant phenotypes of these three mutants, a targeted gene knockout (KO) using homologous recombination was carried out to obtain deletion mutants of *SsSte50* in WT background (Figure 2B). Two independent deletion alleles were obtained and purified. The presence of an amplified fragment within the *SsSte50* gene in WT but not in the two KO mutants, and the presence of the selection marker hygromycin-resistance gene (*HYG*) amplicons in only the two KO mutants confirmed the desired gene replacement (Figure 2C). In the two deletion alleles *Ssste50-4,* and *Ssste50-5*, we observed identical defects as *Ssste50-1*, which do not form sclerotia and grow slower with thick mycelia (Figure 2D-E). Moreover, they could not infect *N. benthamiana* or *A. thaliana* leaves (Figure 2F-I). Since *SsSte50-1* could produce normal amount of OA (Figure 1C), we further tested whether the loss of virulence of in all these *Ssste50* mutants is due to penetration defects. As shown in Figure 2J, these mutants were not able to form compound appressoria at all (Liang et al., 2018). When the *Ssste50* alleles were inoculated onto pre-wounded *N. benthamiana* leaves, they could cause lesions, but the lesion sizes were much smaller compared to WT (Figure 2K-L). Therefore, the virulence defects of *Ssste50* seem to be primarily caused by the failed compound appressoria formation.

In parallel to knocking out *SsSte50*, we tested whether replacement of *Ssste50-1* with full length WT *SsSte50* sequence including its native promoter region via homologous recombination could revert the mutant phenotypes back to WT-like (Figure 3A). The presence of two amplified fragments containing *SsSte50* with the *HYG* marker or *HYG* marker with the downstream fragment (DF) in the complementation strains but not in WT confirmed that the gene replacement was successful (Figure 3B). The two complementation alleles *SsSte50-*C1 and *SsSte50-*C2 formed dark-colored sclerotia and exhibited vegetative growth rates similar to WT (Figure 3C-D). When inoculated on *N. benthamiana* and *A. thaliana* leaves, the *SsSte50* complementation strains caused similar lesion sizes as WT (Figure 3E-H), and they were able to produce compound appressoria (Figure 3I). Thus, the replacement of *Ssste50* in *Ssste50-1* with WT *SsSte50* successfully restored the growth, sclerotia formation, and virulence to the WT phenotypes.

**Figure 3.**
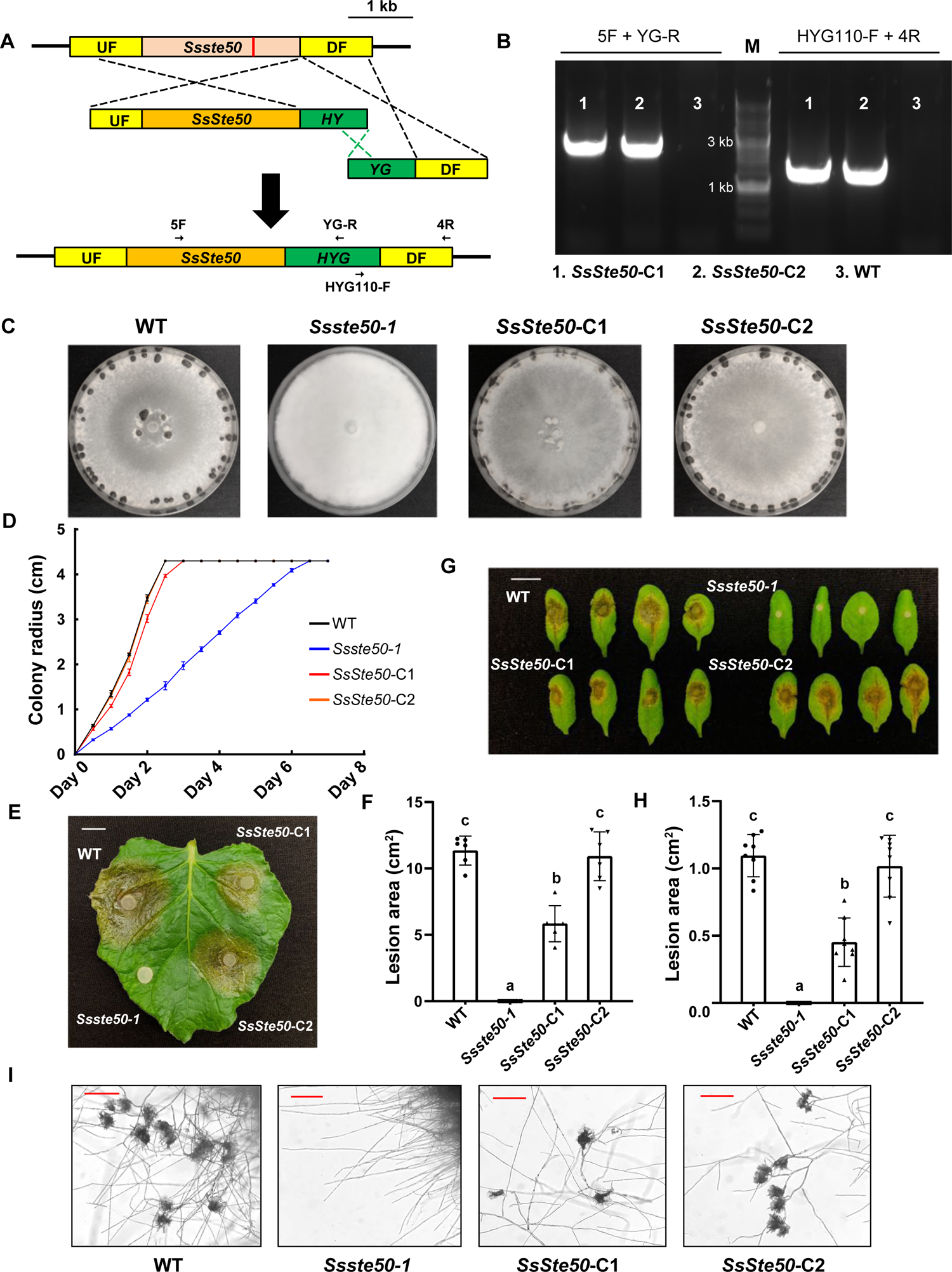
*SsSte50* can complement *Ssste50-1* mutant phenotypes. A. Target gene knock-in (KI) design by homologous recombination. The mutant *Ssste50* gene is shown as a light orange rectangle with the mutation indicated as a red line. The WT *SsSte50* gene and hygromycin-resistance gene *HYG* are shown as orange and green rectangles respectively. *HYG* is split into two halves *HY* and *YG* with overlapping consensus sequence. The primers indicated with arrows are used to genotype the KI transformants. UF: upstream fragment; DF: downstream fragment. B. PCR verification of *SsSte50* gene KI. The genomic DNA from WT *S. sclerotiorum* and two KI strains *SsSte50-*C1, *SsSte50-*C2 were used as PCR templates. The two primer pairs were used to test the insertion of *HYG* gene after *SsSte50* KI. Lane M is the DNA ladder. C. Colony morphology of the indicated genotypes on PDA plates. The pictures were taken at 14 dpi. D. Mycelial growth of the indicated genotypes on PDA plates. The radius was measured every 12 hours until the fungus colonizes the whole plate. Two independent experiments were carried out with similar results. E. Pathogenicity test of the indicated genotypes on detached *N. benthamiana* leaves. The picture in E was taken at 36 hpi. F. Quantification of the lesion areas caused by the indicated *S. sclerotiorum* strains at 36 hpi. Statistical significance is indicated by different letters (*p* < 0.01). Error bars represent means ± SD (*n* = 6). Three independent experiments were carried out with similar results. Bar = 1 cm. G. Pathogenicity test of the indicated genotypes on detached *A. thaliana* Col-0 leaves. The picture in G was taken at 36 hpi. H. Quantification of the lesion areas caused by the indicated *S. sclerotiorum* strains at 36 hpi. Statistical significance is indicated by different letters (*p* < 0.01). Error bars represent means ± SD (*n* = 8). Three independent experiments were carried out with similar results. Bar = 1 cm. I. Compound appressoria formation by the indicated mutants. The mutants were transferred from PDA plates onto glass slides for compound appressoria observation. The pictures were taken at 2 dpi. The Bar = 200 μm.

Taken together, these results from both knockout and complementation experiments confirmed that the mutations in *SsSte50* are responsible for all the developmental and virulence defects observed in the original UV mutants. *SsSte50* gene is thus necessary for normal vegetative growth, sclerotia formation, compound appressoria formation and pathogenicity in host plants.

### A MAP kinase cascade orthologous to the yeast Ste11-Ste7-Kss1/Fus3 in *S. sclerotiorum*, consisting of SsSte11, SsSte7 and Smk1, is similarly involved in development and pathogenicity as SsSte50

SsSte50 being part of a MAPK cascade prompted us to further characterize this putative cascade containing in *S. sclerotiorum*. We first investigated the ortholog of MAPKKK Ste11, which interacts with adaptor Ste50 in yeast (Chen & Thorner, 2007; Hamel et al., 2012). Another mutant M30 identified from our forward genetic screen shared similar mycelial morphology as *Ssste50-1* at initial growth stage (Figure 4A). Interestingly, we noticed that M30 contains a point mutation in the *Ste11* homolog *sscle_03g025710* (Figure 4B), and the mutated serine residue in the protein kinase domain is highly conserved among fungal species (Supplemental Figure S4). The protein structure of sscle_03g025710 kinase domain also shares a high identity with yeast Ste11 (Supplemental Figure S5). *sscle_03g025710* was thus named as *SsSte11* and considered as the top candidate gene for M30. M30 was renamed as *SsSte11-1*.

**Figure 4.**
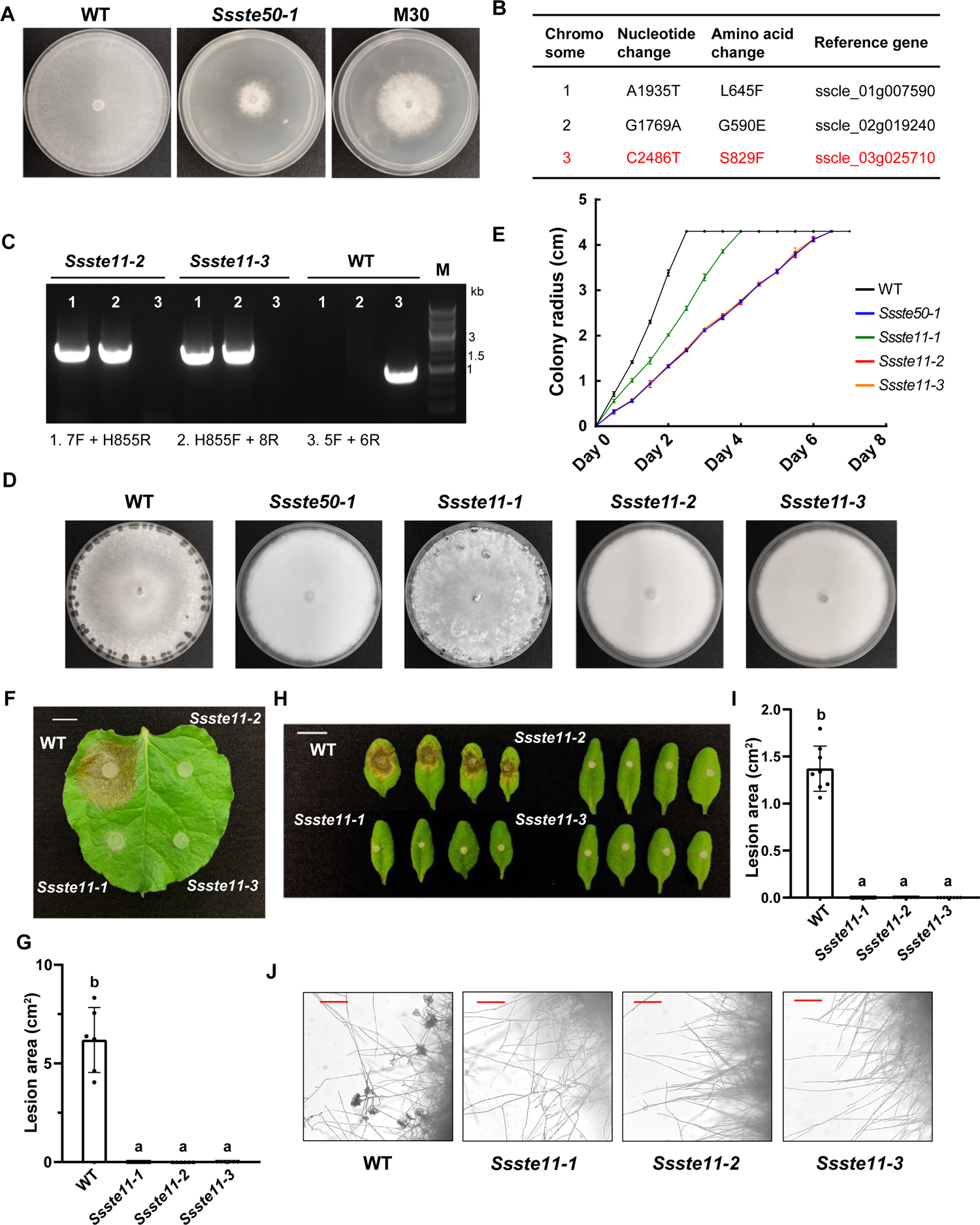
A point mutation in MAPKKK-encoding *SsSte11* accounts for M30 mutant phenotypes and knocking out *SsSte11* phenocopies the *Ssste50* mutants. A. Colony morphology of WT, *Ssste50-1*, and M30 on potato dextrose agar (PDA) plates. The pictures were taken at 60 hpi. B. Candidate mutations of M30 from NGS data analysis. The most likely gene *sscle_03g025710* is highlighted in red. C. PCR verification of *SsSte11* gene KO. The genomic DNA from WT *S. sclerotiorum* and two KO mutants *Ssste11-2*, *Ssste11-3* were used as PCR templates. Primer pairs 1 and 2 were used to test the *HYG* gene insertion, and primer pair 3 was used to test the deletion of *SsSte11*. Lane M is the DNA ladder. D. Colony morphology of WT, *Ssste50-1*, *SsSte11-1*, *Ssste11-*2, and *Ssste11-3* on PDA plates. The pictures were taken at 14 dpi. E. Mycelial growth of the indicated genotypes on PDA plates. The radius was measured every 12 hours until the fungus colonizes the whole plate. Two independent experiments were carried out with similar results. F. Pathogenicity test of the indicated genotypes on detached *N. benthamiana* leaves. The picture in F was taken at 30 hpi. G. Quantification of the lesion areas caused by the indicated *S. sclerotiorum* strains at 30 hpi. Statistical significance is indicated by different letters (*p* < 0.01). Error bars represent means ± SD (*n* = 6). Three independent experiments were carried out with similar results. Bar = 1 cm. H. Pathogenicity test of the indicated genotypes on detached *A. thaliana* Col-0 leaves. The picture in H was taken at 36 hpi. I. Quantification of the lesion areas caused by the indicated *S. sclerotiorum* strains at 36 hpi. Statistical significance is indicated by different letters (*p* < 0.01). Error bars represent means ± SD (*n* = 8). Three independent experiments were carried out with similar results. Bar = 1 cm. J. Compound appressoria formation by the indicated mutants. The mutants were transferred from PDA plates onto glass slides for compound appressoria observation. The pictures were taken at 2 dpi. The Bar = 200 μm.

To test whether the mutation in *SsSte11* is responsible for its developmental and pathogenicity defects, deletion mutants of *SsSte11* in WT background were generated (Figure 4C). The two purified deletion alleles *Ssste11-2* and *Ssste11-3* exhibited similar sclerotia formation defects as *Ssste50-1* (Figure 4D). *SsSte11-1* showed higher vegetative growth rate compared to the *Ssste11* KO mutants (Figure 4E). We reasoned that since *SsSte11-1* carries a single amino acid change in SsSte11, its protein is likely still partially functional to support sclerotia formation and vegetative growth, though not to the WT level. Lastly, the *SsSte11-1* and the two *Ssste11* deletion alleles could not infect *N. benthamiana* or *A. thaliana* leaves (Figure 4F-I). The lack of compound appressoria formation suggests that these mutants are unable to penetrate the plant epithelial layer to initiate infection (Figure 4H).

As SsSte11 is a MAPKKK, we next sought to test the downstream MAPKK and MAPK. Since MAPK cascades are highly conserved in fungi, *SsSte7* (*sscle_09g069880*) and *Smk1* (*sscle_12g090900*) seem to be the orthologous candidates according to previous phylogeny analyses and our own protein BLAST and structural examinations (Supplemental Figure S6-S7; Hamel et al., 2012; Hegedus et al., 2016). When pure deletion alleles of *SsSte7* and *Smk1* were generated, they were incapable of forming sclerotia (Figure 5A-C). They displayed similar vegetative growth and inability to infect host plant leaves (Figure 5D-H). Like *Ssste50-1*, they were unable to form compound appressoria (Figure 5I). Such identical phenotypes among *Ssste50*, *Ssste11*, *Ssste7*, and *smk1* mutants indicate a clear MAPK cascade formed by their encoded kinases or kinase adaptor regulating *S. sclerotiorum* development and virulence.

**Figure 5.**
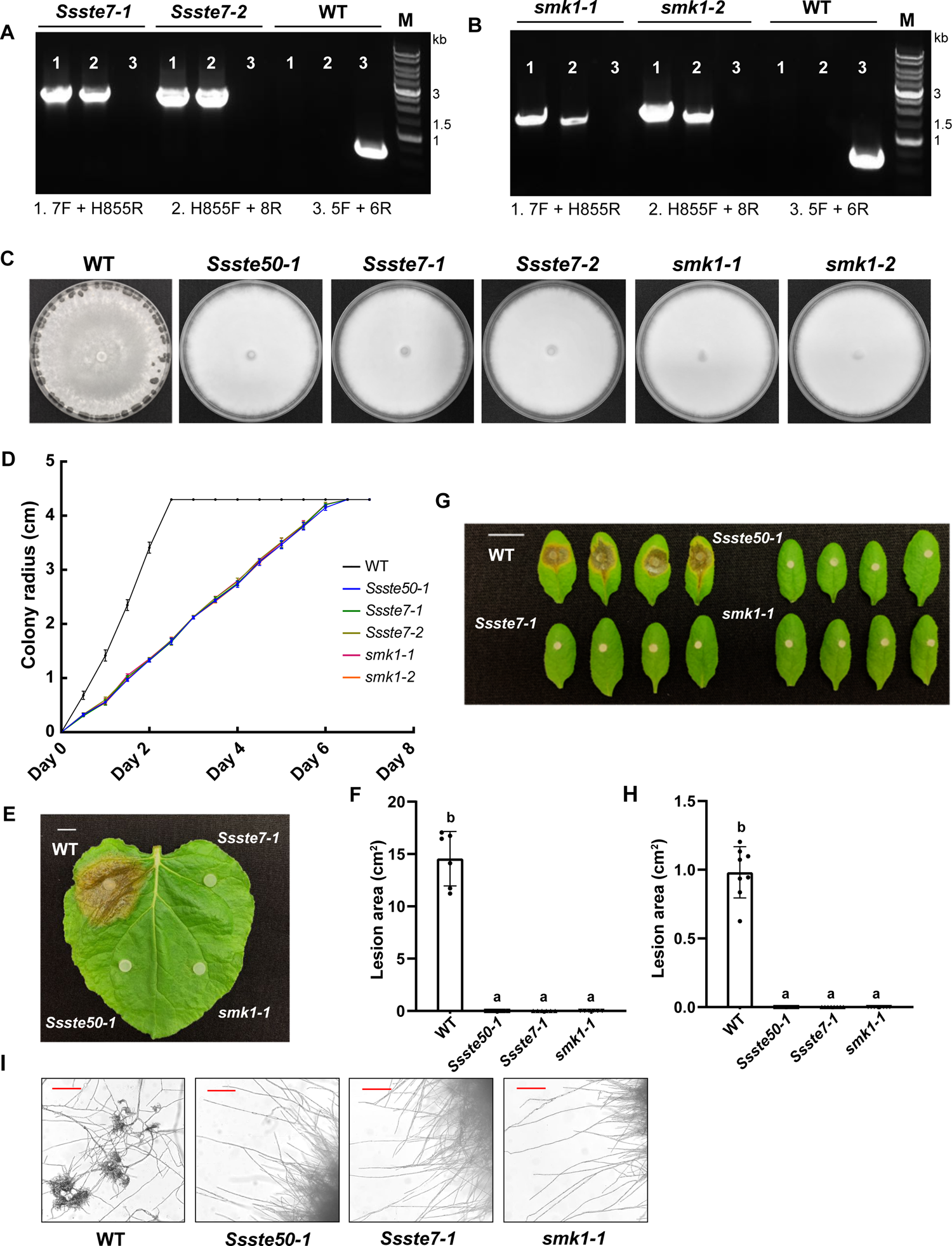
knocking out MAPKK-encoding *SsSte7* or MAPK-encoding *Smk1* phenocopies *Ssste50* mutants. A and B. PCR verification of *SsSte7* and *Smk1* gene KOs. The genomic DNA from WT *S. sclerotiorum* and KO mutants *Ssste7-1*, *Ssste7-2*, *smk1-1*, *smk1-2* were used as PCR templates. Primer pairs 1 and 2 were used to test the *HYG* gene insertion, and primer pair 3 was used to test the target gene deletion. Lane M shows the DNA ladder. C. Colony morphology of the indicated genotypes on PDA plates. The pictures were taken at 14 dpi. D. Mycelial growth of the indicated genotypes on PDA plates. The radius was measured every 12 hours until the fungus colonizes the whole plate. Two independent experiments were carried out with similar results. E. Pathogenicity test of the indicated genotypes on detached *N. benthamiana* leaves. The picture in E was taken at 40 hpi. F. Quantification of the lesion areas caused by the indicated *S. sclerotiorum* strains at 40 hpi. Statistical significance is indicated by different letters (*p* < 0.01). Error bars represent means ± SD (*n* = 6). Three independent experiments were carried out with similar results. Bar = 1 cm. G. Pathogenicity test of the indicated genotypes on detached *A. thaliana* Col-0 leaves. The picture in G was taken at 36 hpi. H. Quantification of the lesion areas caused by the indicated *S. sclerotiorum* strains at 36 hpi. Statistical significance is indicated by different letters (*p* < 0.01). Error bars represent means ± SD (*n* = 8). Three independent experiments were carried out with similar results. Bar = 1 cm. I. Compound appressoria formation by the indicated mutants. The mutants were transferred from PDA plates onto glass slides for compound appressoria observation. The pictures were taken at 2 dpi. The Bar = 200 μm.

### The putative transcription factor SsSte12 activated by MAPK is required for compound appressoria development and host penetration

Ste12 is the major transcription factor activated by the pheromone response MAPK cascade in yeast, which regulates the mating process (Fields et al., 1988; Hamel et al., 2012). To investigate the contribution of its homolog *SsSte12* (*sscle_06g051560*) in development and virulence in *S. sclerotiorum*, we deleted *SsSte12* by gene KO (Figure 6A). Surprisingly, the two purified deletion alleles, *Ssste12-1* and *Ssste12-2,* formed sclerotia similar with WT (Figure 6B). In contrast to the upstream MAPK components, they grew slightly slower than WT but faster than *Ssste50-1* (Figure 6C). Moreover, when infected on host leaves, *Ssste12* mutants showed smaller lesion sizes compared with WT (Figure 6D-G). Interestingly, they developed distorted, expansive appressoria (Figure 6H). These appressoria are likely less able to penetrate, thereby contributing to the decreased lesion size. Indeed, inoculation of the *Ssste12* mutants onto pre-wounded *N. benthamiana* leaves produced lesions comparable to WT (Figure 6I-J). These results suggest that the SsSte12, unlike the MAPK components, is not necessary for sclerotia formation. However, it is required for normal compound appressoria development during host penetration. The reduction of defects from MAPK cascade mutants observed in *Ssste12* suggests that there likely are multiple transcription factors (TFs) downstream of the cascade, SsSte12 is only in charge of a portion of the events downstream.

**Figure 6.**
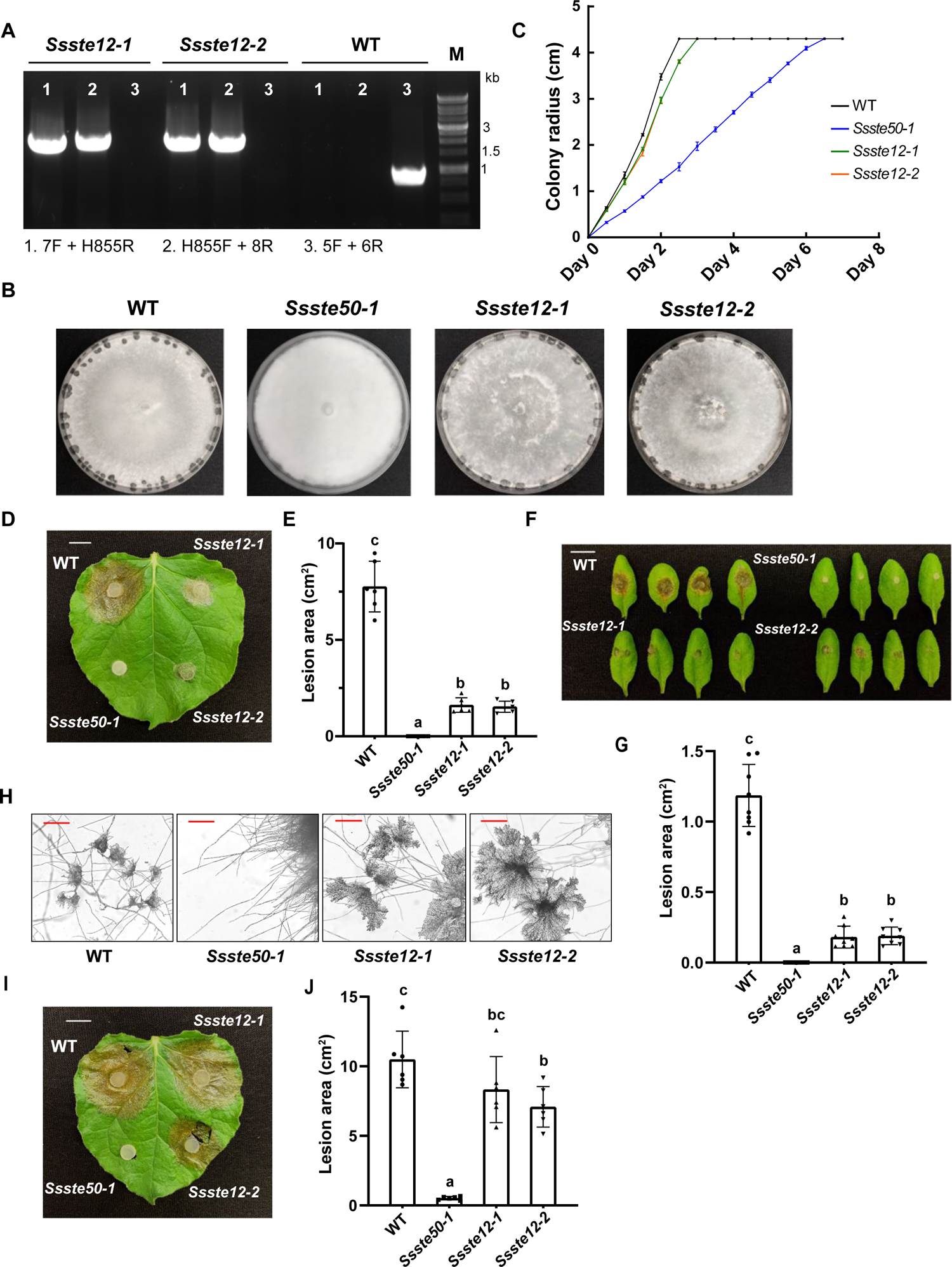
Transcription factor SsSte12 is required for compound appressoria formation and host penetration. A. PCR verification of *SsSte12* gene KO. The genomic DNA from WT *S. sclerotiorum* and KO mutants *Ssste12-1*, *Ssste12-2* were used as PCR templates. Primer pairs 1 and 2 were used to test the *HYG* gene insertion, and primer pair 3 was used to test the deletion of *SsSte12*. Lane M shows the DNA ladder. B. Colony morphology of the indicated genotypes on PDA plates. The pictures were taken at 14 dpi. C. Mycelial growth of the indicated genotypes on PDA plates. The radius was measured every 12 hours until the fungus colonizes the whole plate. Two independent experiments were carried out with similar results. D. Pathogenicity test of the indicated genotypes on detached *N. benthamiana* leaves. The picture in D was taken at 30 hpi. E. Quantification of the lesion areas caused by the indicated *S. sclerotiorum* strains at 30 hpi. Statistical significance is indicated by different letters (*p* < 0.01). Error bars represent means ± SD (*n* = 6). Three independent experiments were carried out with similar results. Bar = 1 cm. F. Pathogenicity test of the indicated genotypes on detached *A. thaliana* Col-0 leaves. The picture in F was taken at 36 hpi. G. Quantification of the lesion areas caused by the indicated *S. sclerotiorum* strains at 36 hpi. Statistical significance is indicated by different letters (*p* < 0.01). Error bars represent means ± SD (*n* = 8). Three independent experiments were carried out with similar results. Bar = 1 cm. H. Compound appressoria formation by the indicated mutants. The mutants were transferred from PDA plates onto glass slides for compound appressoria observation. The pictures were taken at 2 dpi. The Bar = 200 μm. I. Pathogenicity test of the indicated genotypes on detached *N. benthamiana* leaves with pre-inoculation wounding. The picture in I was taken at 30 hpi. J. Quantification of the lesion areas caused by the indicated *S. sclerotiorum* strains at 30 hpi. Statistical significance is indicated by different letters (*p* < 0.01). Error bars represent means ± SD (*n* = 6). Three independent experiments were carried out with similar results. Bar = 1 cm.

### SsSte50 is required for *S. sclerotiorum* hyphal fusion

Hyphal fusion is ubiquitous in filamentous fungi, playing an essential role in intra-hyphal communication, nutrient exchange, and mating processes (Glass et al., 2004; Fischer & Glass, 2019). Although the molecular mechanisms behind hyphal fusion are not fully understood, components from the pheromone response MAPK pathway including Ste50 and Ste11 are known to be required for proper communication and fusion between cells in *S. cerevisiae* and *Neurospora crassa* (Glass et al., 2004; Dettmann et al., 2014). To interrogate whether the MAPK cascade also plays a role in *S. sclerotiorum* hyphal fusion, we designed a quick and efficient approach taking advantage of different *S. sclerotiorum* mutants generated from our UV mutagenesis (Figure 7A). The pink-sclerotia mutant *Sssmr1-1* (Figure 7B, Xu et al., 2022) without fusion defects was co-inoculated with another non-sclerotial forming mutant on PDA plates. If the black sclerotia can form in the middle, such mutant is considered without fusion defects since the black sclerotia were produced, indicating successful fusion, exchange of nuclei and complementation of the mutant phenotypes (Figure 7A). When *Sssmr1-1* was co-inoculated with a non-sclerotial forming mutant R240 (the causal gene for R240 is not identified yet) (Figure 7B, Xu et al., 2022), several black sclerotia can be observed in the middle, suggesting that the mutated gene in R240 is not required for hyphal fusion (Figure 7C). However, when tested with *Ssste50-1* and *Ssste50-2*, no black sclerotia can be detected and a few pink sclerotia were formed at the borderline, indicative of fusion defects. Interestingly, *Ssste50-3* with a more downstream mutation (Figure 2A) retained partial fusion ability as the sclerotia melanisation occurred (Figure 7C). This agrees with our previous results that it is likely a partial loss-of-function allele. Collectively, these data demonstrate that SsSte50 is indispensable for proper fusion between *S. sclerotiorum* hyphal cells, and the downstream Ras-associating domain of SsSte50 may be less important for such processes.

**Figure 7.**
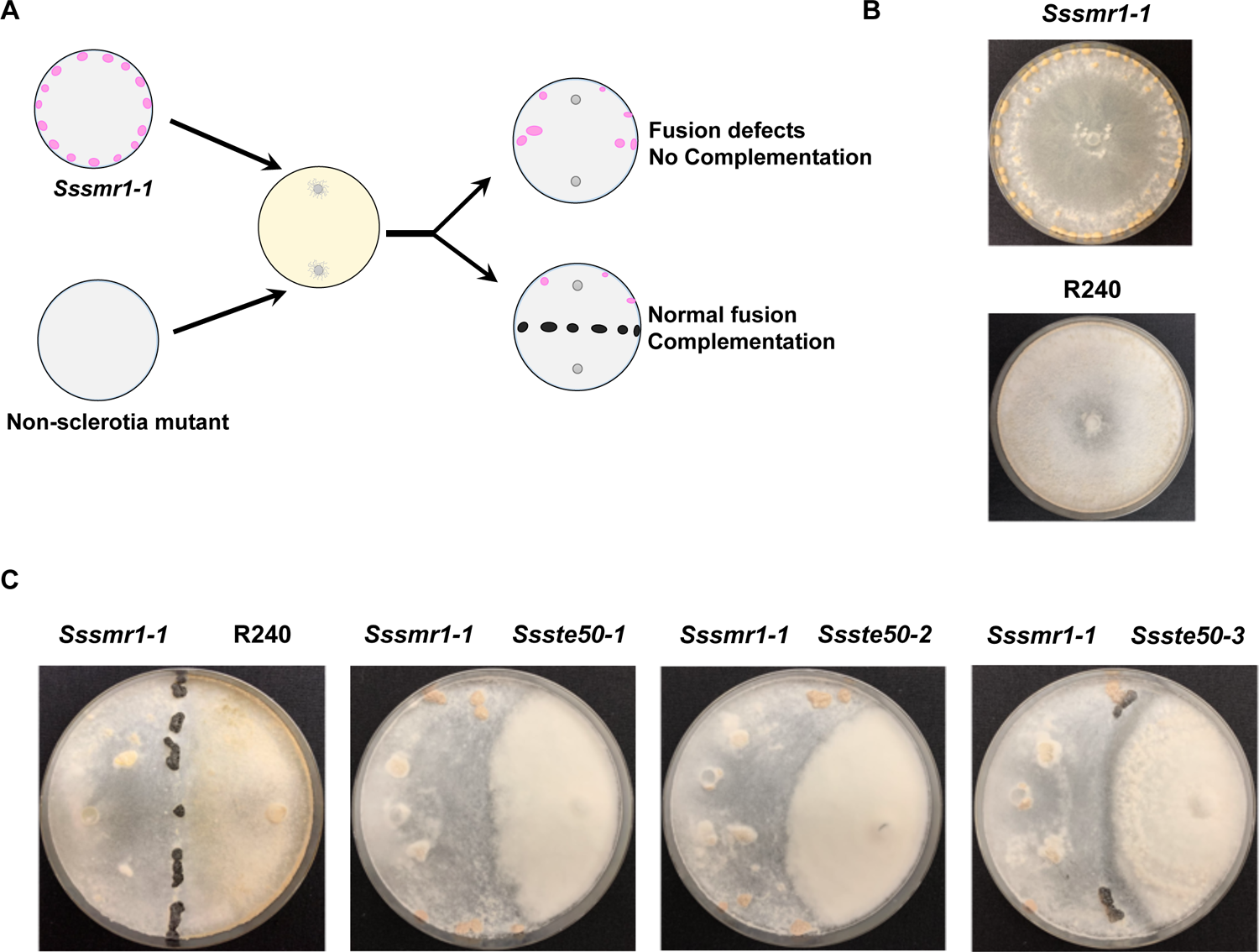
SsSte50 is indispensable for hyphal fusion in *S. sclerotiorum*. A. A diagram depicting the mutant pairing experiment designed for testing hyphal fusion ability of *S. sclerotiorum* mutants. B. Colony morphology of *Sssmr1-1* that forms pink sclerotia and R240 that does not form sclerotia on PDA plates. The pictures were taken at 14 dpi. C. Hyphal fusion test of *Ssste50-1*, *Ssste50-2*, and *Ssste50-3* with *Sssmr1-1*. R240 was used as a positive control. As the *Ssste50* mutans grow slower, Ssste50*-1*, *Ssste50-2*, and *Ssste50-3* were inoculated on PDA plates 2 days prior to *Sssmr1-1* inoculation. The pictures were taken at 14 dpi.

### HIGS of SsSte50 in N. benthamiana compromised S. sclerotiorum virulence

In recent years, HIGS has become a valuable approach to control phytopathogenic fungi. It is based on engineering plants to express double-stranded RNAs (dsRNAs) that target important pathogen mRNAs for degradation (Hua et al., 2018). It has been successfully applied to multiple plant species for disease control (Tinoco *et al*., 2010; Nowara *et al*., 2010; Zhang *et al*., 2016). In *S. sclerotiorum*, HIGS was designed to target known virulence genes (McCaghey *et al*., 2021; Rana *et al*., 2021; Wytinck *et al*., 2022). Since the MAPK cascade consisting of SsSte50-SsSte11-SsSte7-Smk1 is essential for pathogenesis of *S. sclerotiorum*, we tested if it can be a HIGS target for disease control. When the *SsSte50* hairpin RNAi construct for HIGS was expressed in *N. benthamiana* leaves, a dramatic reduction in necrotic lesion size caused by *S. sclerotiorum* was observed compared to the empty vector (EV)-treated control (Figure 8). This successful trans-species RNAi targeting *SsSte50* suggests that this MAPK cascade can serve as an excellent HIGS target to prevent *S. sclerotiorum* infection.

**Figure 8.**
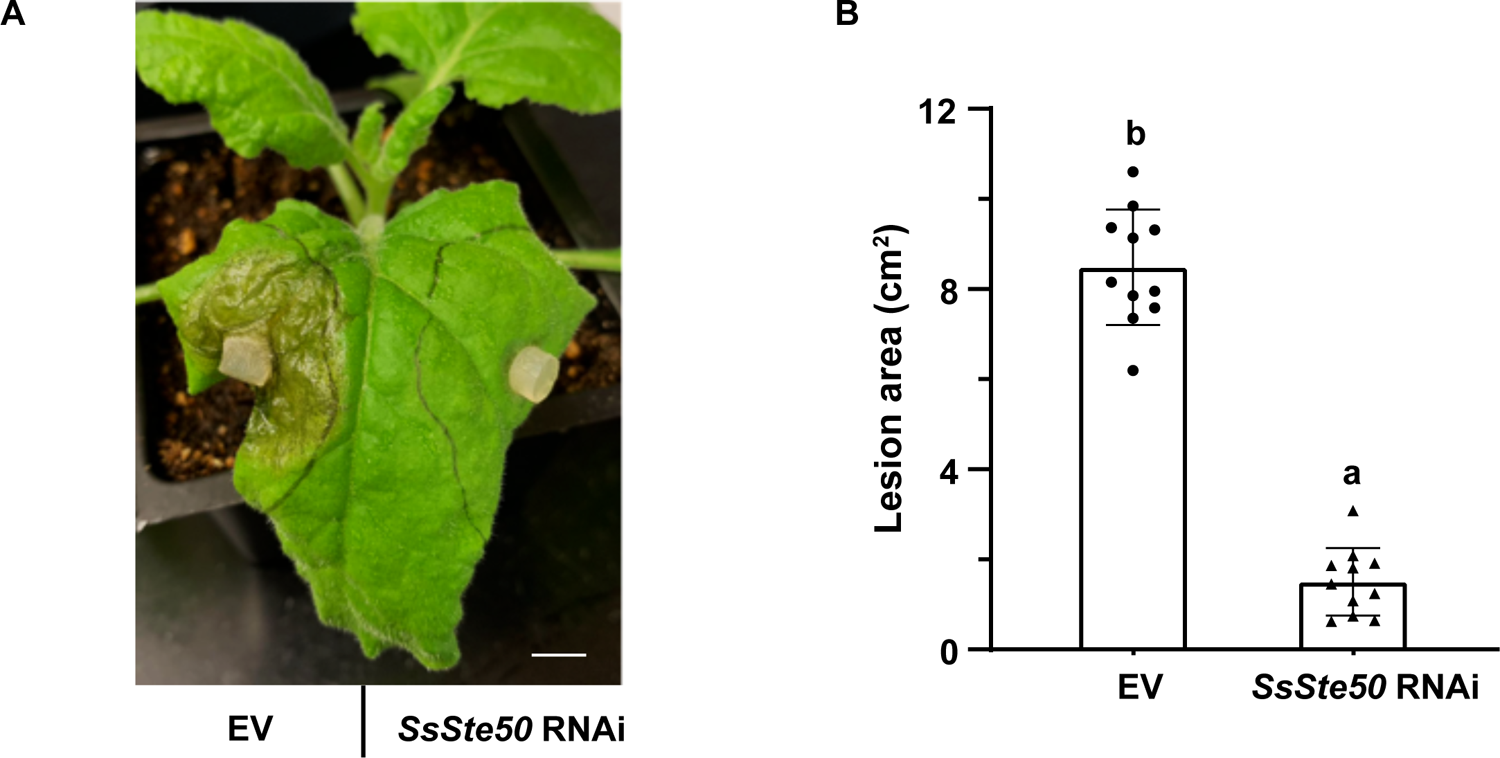
HIGS of *SsSte50* compromises *S. sclerotiorum* virulence in *N. benthamiana*. A. Pathogenicity test of WT *S. sclerotiorum* strain on *N. benthamiana* leaves expressing EV (left) and *SsSte50* hairpin RNAi (right) constructs. The picture was taken at 36 hpi. B. Quantification of the lesion areas at 36 hpi. Statistical analysis was carried out with one-way ANOVA followed by Tukey’s post hoc test. Statistical significance is indicated by different letters (*p* < 0.01). Error bars represent means ± SD (*n* = 11). Three independent experiments were carried out with similar results. Bar = 1 cm.

## Discussion

Although MAPK cascades are known signaling masters in eukaryotes, they have not been well studied in *S. sclerotiorum*. In this study, we characterized a highly conserved cascade consisting of SsSte50-SsSte11-SsSte7-Smr1, which broadly regulates vegetative growth, sclerotia formation, compound appressoria formation and pathogenicity. The putative downstream transcription factor SsSte12, however, is mainly required for compound appressoria development and host penetration (Figure 9). The parallel pheromone response MAPK signaling pathway in *S. cerevisiae* and *N. crassa* plays an essential role in hyphal cell fusion (Glass et al., 2004; Dettmann et al., 2014). Likewise, SsSte50 is also indispensable for hyphal fusion in *S. sclerotiorum* (Figure 7).

**Figure 9.**
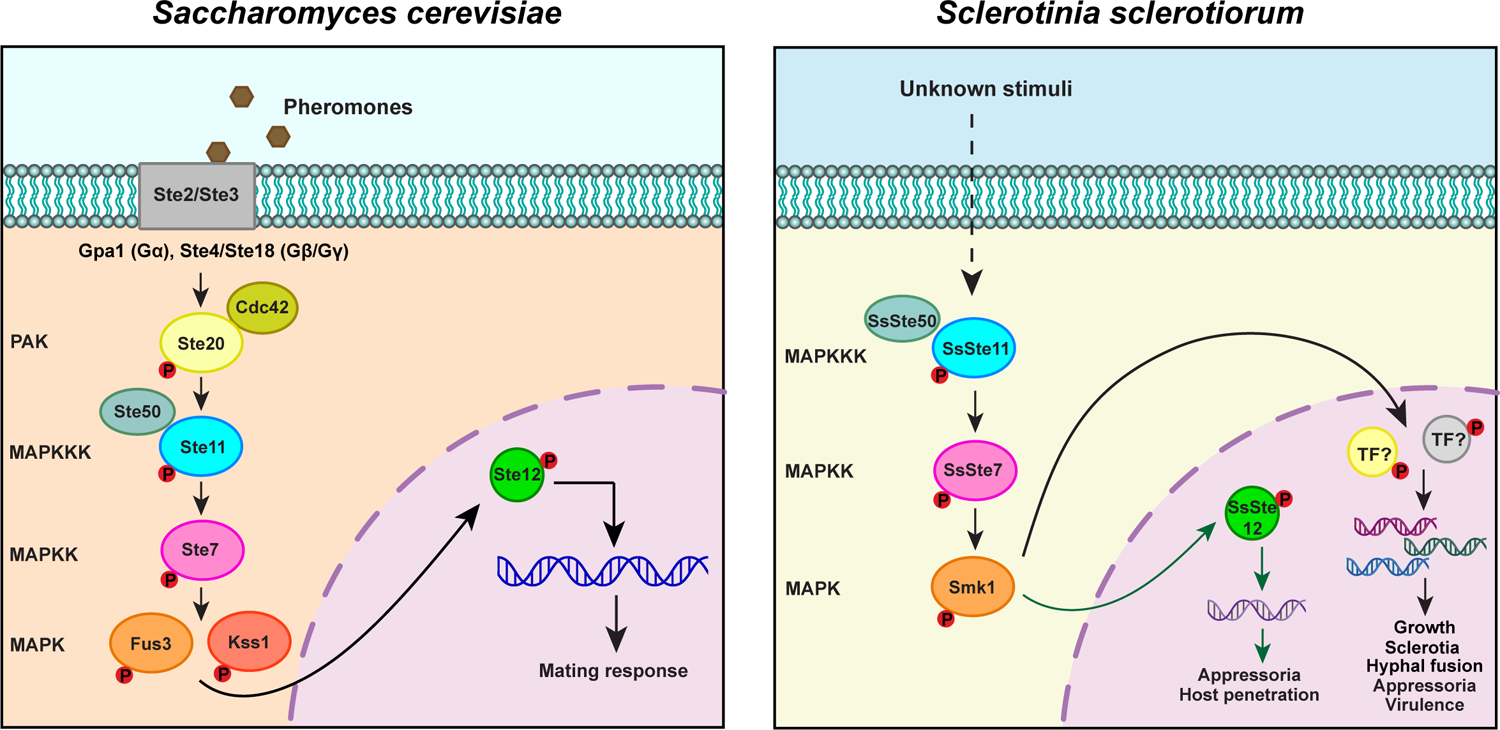
Comparable models of the pheromone response MAP kinase signaling pathway in *S. cerevisiae* and *S. sclerotiorum*. In *S. cerevisiae*, G-protein-coupled receptors (GPCRs) Ste2 or Ste3 recognize the cognate pheromone signals, which activates the G protein module consisting of Gpa1 (Gα), Ste4/Ste18 (Gβ/Gγ). The released Gβ Ste4 then activates a p21-activated protein (PAK) kinase Ste20 possibly with a Rho protein Cdc42. The sequential activation of the MAPK cascade components is initiated by Ste20, and the partially redundant downstream MAPK Fus3 and Kss1 mostly phosphorylate transcription factor Ste12 for regulating the mating process. In parallel in *S. sclerotiorum*, upon detection of unknown upstream stimuli, a MAPK cascade consisting of the adaptor protein SsSte50, MAPKKK SsSte11, MAPKK SsSte7, MAPK Smk1 is activated. The phosphorylated Smk1 then activates the transcription factor (TF) SsSte12 and other yet-to-be identified TFs (TF?) to regulate vegetative growth, sclerotia formation, compound appressoria formation, hyphal fusion, and virulence.

The well-established pheromone response pathway in yeast *S. cerevisiae* (Figure 9) starts with the recognition of pheromone signals by G-protein-coupled receptors (GPCRs) Ste2 or Ste3. The sequential phosphorylation of the MAPK cascade components occurs after activation of a G protein module and the p21-activated protein (PAK) kinase Ste20. The partially redundant downstream MAPK Fus3 and Kss1 then phosphorylate transcription factor Ste12 for regulating the mating process (Neiman and Herskowitz, 1994; Sabbagh et al., 2001; Zhao et al., 2007; Hamel et al., 2012). Homology and phylogenetic analyses of this Ste50-Ste11-Ste7-Kss1/Fus3 MAPK cascade revealed its high conservation among fungal species (Zhao et al., 2007; Hamel et al., 2012; Hegedus et al., 2016). Likewise, our protein sequence and structure examinations confirmed such identity in phytopathogenic fungi including *S. sclerotiorum* (Supplemental Figure S3-S7). Over the past two decades, elaborate characterization of this pheromone response MAPK cascade has been carried out in some phytopathogenic fungal species including *M. oryzae* (Zhao et al., 2005; Park et al., 2006), corn smut *Ustilago maydis* (MuLller et al., 2003), poplar anthracnose fungus *Colletotrichum gloeosporioides* (Wang et al., 2021) and *B. cinerea* (Schamber et al., 2010). All these studies indicate a common role of this MAPK cascade in fungal growth and host penetration, mostly through compound appressoria formation, which is consistent with our *S. sclerotiorum* data. Interestingly, the cascade is also required for hyphal fusion and sclerotia formation in *S. sclerotiorum*, which is a unique developmental feature of this fungal family. Since proper sclerotia formation is a prerequisite for entering the sexual cycle (Xia et al., 2020), activation of the MAPK signaling pathway may also be initiated by certain pheromone signals in *S. sclerotiorum* as with in *S. cerevisiae*. Future identification and investigation of the pheromones and their cognate receptors will help to define the upstream events of this MAPK cascade.

The existence of a Ras-associating (RA) domain in most fungal Ste50 and Ste11 homologs (Supplemental Figure S3-S4) and the severe defects observed in *Ssste50-3* allele missing the RA domain (Figure 1-2) suggest that certain Ras family member is required for the MAPK cascade activation. Ras proteins are small GTPases that regulate many signal transduction pathways involved in development, stress responses, and proliferation in fungi (Dautt-Castro et al., 2021). In *S. cerevisiae*, a Rho protein Cdc42 from Ras superfamily associates with the kinase Ste20 and serves as the upstream regulator of the MAPK cascade (Figure 9, Lamson et al., 2002). Cdc42 was also detected to interact with Ste50 (Truckses et al., 2006), confirming the importance of Ras proteins in the pheromone response MAPK signaling. However, although Cdc42 interacts with Mst50 (Ste50 homolog) in *M. oryzae*, it is dispensable for appressoria formation and host infection, indicating Cdc42 is less likely to act in the Mst50 pathway (Park et al., 2006; Zhao et al., 2007). Similar observation was also reported in *B. cinerea* (Schamber et al., 2010; Kokkelink et al., 2011). Given the higher amino acid similarity of the RA domains of SsSte50 and SsSte11 with the *B. cinerea* and *M. oryzae* homologs in contrast to the *S. cerevisiae* homologs (Supplemental Figure S3-S4), Cdc42 is less likely to be the corresponding Ras in *S. sclerotiorum* either. Interestingly, Mst50 and Ste11 also interact with Ras2 in *M. oryzae*. Although lethality of the *ras2* KO mutants hinders the epistasis analysis, expression of an auto-active *Ras2* in WT but not the MAPK mutants stimulated appressoria formation, suggestive of its potential role in the Mst50 pathway (Park et al., 2006). Similar requirement of Ras2 in *U. maydis* MAPK cascade was also reported by using a dominant active *ras2* allele (Lee and Kronstad, 2002). Likewise, complete deletion of *Ras2* gene is lethal in *S. sclerotiorum* (Xu et al, unpublished data from our lab). These data indicate a master role of Ras2 GTPase in fungal signaling. However, whether Ras2 is the Ste50 and Ste7 interactor for MAPK signaling in *S. sclerotiorum* and other phytopathogenic fungi requires further investigations.

Besides the pheromone response pathway, the other two well-conserved high osmolarity and cell wall integrity MAPK cascades serve to alleviate environmental stressors in *S. cerevisae* (Posas & Saito, 1997; Levin, 2011; Hamel et al., 2012). In phytopathogenic fungi, however, these cascades can also be involved in pathogenicity (Xu et al., 1998; Rui and Hahn, 2007; Bashi et al., 2016). For example, knocking out *Sak1* in the *B. cinerea* high osmolarity pathway abolished compound appressoria formation required for host entry (Segmüller et al., 2007). Similarly in *S. sclerotiorum*, disruption of the osmolarity responsive MAPKKK Ssos4 led to reduced sclerotia formation and virulence (Li et al., 2021). Deletion of the cell wall integrity MAPK cascade components not only led to decreased cell wall integrity, but also decreased sclerotia formation and virulence (Bashi et al., 2016; Cong et al., 2022). In contrast, depending on the dysregulated pathway, *S. sclerotiorum* exhibited differing mycelial morphologies, suggesting that downstream components of the pheromone response, high osmolarity and cell wall integrity MAPK pathways likely function independently with some overlap (Li et al., 2021; Cong et al., 2022). However, the details of crosstalk among the three MAPK signaling pathways in regulating *S. sclerotiorum* development and virulence require further careful analysis.

The homologs of yeast pheromone response pathway transcription factor Ste12 are involved in pathogenesis of *B. cinerea* and *S. sclerotiorum* (Schamber et al., 2010; Xu et al., 2018). Consistently, in this study, deletion of *SsSte12* greatly attenuated virulence on intact leaves due to abnormal appressoria formation. Unlike the *Ssste50* KO mutants, *Ssste12* exhibited normal growth and sclerotia formation, and caused similar necrotic lesions on pre-wounded leaves as with WT *S. sclerotiorum*. Thus, there are likely other unknown transcription factors that regulate growth, sclerotia and compound appressoria formation downstream of the MAPK Smk1. For instance, along with Ste12, the transcription factor Tec1 is required for filamentous growth in *S. cerevisae* and *Candida albicans* (Hamel et al., 2012; Schweizer et al., 2000). However, close homologs of Tec1 cannot be identified by protein BLAST in *S. sclerotiorum*. Interestingly, through a screen for phosphorylated transcription factor by the MAPK Pmk1 in *M. oryzae*, Sfl1 was identified, and deletion of *Sfl1* showed similar appressoria defects as the MAPK mutants (Li et al., 2011). Besides, *Gas1* and *Gas2* encoding certain small proteins are regulated by Pmk1, and they were required for host penetration (Xue et al., 2002). Conceivably, homologs of Sfl1, Gas1, and Gas2 may also contribute to the Smr1 MAPK signaling pathway in *S. sclerotiorum*. However, downstream factors that regulate vegetative growth and sclerotia formation in *S. sclerotiorum* remain largely unknown.

HIGS technology has been developed rapidly in recent years as an effective approach to protect plants from pathogen or pest infections. It allows the production of designed dsRNAs in host plants that silence pathogenicity-related genes in invaders. There are currently over 170 reports of successful HIGS attempts (Koch and Wassenegger, 2021). However, as a less studied important soilborne fungal pathogen, good HIGS targets for *S. sclerotiorum* are limited. The MAPK cascade we characterized here is indispensable for host penetration and full virulence of *S. sclerotiorum*. Therefore, components from such cascade are potential targets of HIGS for controlling *S. sclerotiorum* diseases in plants. Indeed, HIGS of the MAPK adaptor gene *SsSte50* in tobacco *N. benthamiana* plants largely reduced necrotic lesions caused by *S. sclerotiorum*. Since SsSte50 and SsSte11/SsSte7/Smr1 are activated sequentially during pathogenesis of *S. sclerotiorum*, HIGS of these MAPK cascade genes is most likely to be similarly effective in *N. benthamiana*. Moreover, these MAPK components are ideal HIGS targets for controlling white mold/stem rot diseases in other plant hosts including major crops like canola. In a broader sense, as the MAPK cascade is highly conserved in phytopathogenic fungi for virulence (Hamel et al., 2012), it may also be potentially utilized for controlling other fungal diseases.

## Materials and Methods

### Fungal strains and culture conditions

The wild-type (WT) strain *S. sclerotiorum* 1980 and all the mutant strains generated in the background were maintained on potato dextrose agar (PDA, Shanghai Bio-way technology) at room temperature and stored on PDA slants at 4 °C or as sclerotia. *S. sclerotiorum* KO and KI mutants were screened and purified on PDA supplemented with hygromycin B (Sigma) at a final concentration of 50 μg/ml.

### Oxalic acid (OA) production analysis

To assay the OA production, bromophenol blue (VWR Life Science) was added to PDA at a final concentration of 50 mg/L. The Petri dish used is of 60×15 mm size (SARSTEDT).

### *S. sclerotiorum* genomic DNA extraction and NGS analysis

Extraction of *S. sclerotiorum* genomic DNA for NGS and the NGS data analysis were carried out as described previously (Xu et al., 2022). For NGS data analysis, the sequence reads were mapped to WT strain *S. sclerotiorum* 1980 genome as the reference by BWA-MEM (Li 2013). SAMtools was applied to identify the mutation sites (Li et al. 2009). The annotation of each mutation was done by germline short variant discovery (SNPs + Indels) based on GATK best practices (Van der Auwera et al. 2013).

### Target gene knockout (KO) and transgene complementation (knock-in, KI)

The split marker approach used to make *SsSte50*, *SsSte11*, *SsSte7*, *Smk1*, and *SsSte12* gene replacement cassettes and generation of the KO mutants were described previously (Xu et al., 2022). In brief, the gene replacement cassette consisting of the gene upstream sequence, hygromycin-resistance gene *HYG*, and the gene downstream sequence was transformed into WT *S. sclerotiorum* protoplasts. The positive transformants were screened from the PDA media supplemented with hygromycin B. The KO mutants were further purified by the tip mycelia transfer for several rounds until no WT band can be amplified by PCR using 5F and 6R primer pair. Transgene complementation was done with a similar KI method. The gene replacement cassettes for *SsSte50* include two separate fragments. Fragment 1 consists of *SsSte50* upstream sequence and WT *SsSte50*, as well as a half of *HYG* (*HY*) sequence. Fragment 2 consists of the other half of *HYG* (*YG*) sequence and *SsSte50* downstream sequence. *HY* and *YG* have around 150 bp overlap. The two fragments were co-transformed into Z-2 *Ssste50-1* protoplasts. Subsequent steps including selection of positive transformants, and KI mutant purification were the same with KO methods. All the primers used are listed in Supplemental Table S1.

### *S. sclerotiorum* growth rate determination and colony morphology observation

All the *S. sclerotiorum* WT and mutant strains were grown on PDA media for 2-3 days. The mycelial agar disks made by using a sterile hole punch were then transferred from the colony margin to the center of new PDA plates (92×16 mm, SARSTEDT). The colony radius was measured every 12 hours until the mycelia reach the edge of the paltes at room temperature. The colony morphology pictures were taken 14 days post inoculation for sclerotia observation.

### Plant infection assay

Fresh mycelial plugs (2 mm or 5 mm in diameter) of 2-day-old fresh cultures were inoculated on detached leaves of *A. thaliana* ecotype Col-0 or *N. benthamiana* with or without mechanical wounding. The leaves were placed on moistened paper towels in trays covered with lids, which were then incubated in a growth chamber (23 °C; 16-hrs light/8-hrs dark). The lesion areas were quantified using ImageJ (https://imagej.nih.gov/ij/).

### Compound appressoria observation

Fresh mycelial plugs (5 mm in diameter) taken from the colony margin were placed onto the glass slides on moistened paper towels and incubated at room temperature for 2 days. The formation of compound appressoria was visualized by a ZEISS light microscope.

### *S. sclerotiorum* hyphal fusion test

Fresh mycelial plugs (5 mm in diameter) from the pink sclerotia-forming mutant *Sssmr1-1* (Xu et al., 2022) and another non-sclerotia-forming mutant were placed onto opposite sides of a new PDA plate (Figure 7A). The sclerotia formations were recorded in the middle of the plate after 14 days.

### RNAi-mediated HIGS construct design and transient expression in *N. benthamiana*

To make the HIGS construct for RNAi, 465 bp sense fragment from the coding sequence of *SsSte50* (S) was fused with an intron 3 fragment from *A. thaliana* malate synthase gene (ms-i3; AT5G03860) as described previously (Tinoco *et al*., 2010) by using double-joint PCR. The fused fragment was cloned into a plant expression vector pCambia1300 to generate the intermediate construct pCambia1300-S-i3. Then the 465 bp antisense fragment was ligated into pCambia1300-S-i3 vector to create the final pCambia1300-*SsSte50*-RNAi construct. The primers used for making the HIGS construct were listed in Supplemental Table S1.

For transient expression in *N. benthamiana*, the binary pCambia1300-*SsSte50*-RNAi construct was transformed into *Agrobacterium tumefaciens* GV3101 through electroporation. The agrobacteria harboring empty vector (EV) pCambia1300 construct or the *SsSte50* RNAi construct were infiltrated into four-week-old *N. benthamiana* leaves at OD_600_ = 0.8 as described previously (Wu *et al*., 2022). The plants were kept in dark for 3 days after agrobacteria infiltration to induce the expression of the RNAi construct. After that, the infiltrated *N. benthamiana* leaves were inoculated with WT *S. sclerotiorum* mycelia to test for disease progression.

### Statistical analysis

Statistical analysis was carried out with one-way ANOVA followed by Tukey’s post hoc test. The Scheffé multiple comparison was applied to test correction. Statistical significance was indicated with different letters. *p* values and sample numbers (*n*) were elaborated in figure legends.

---

**Supplemental Figure S1.**
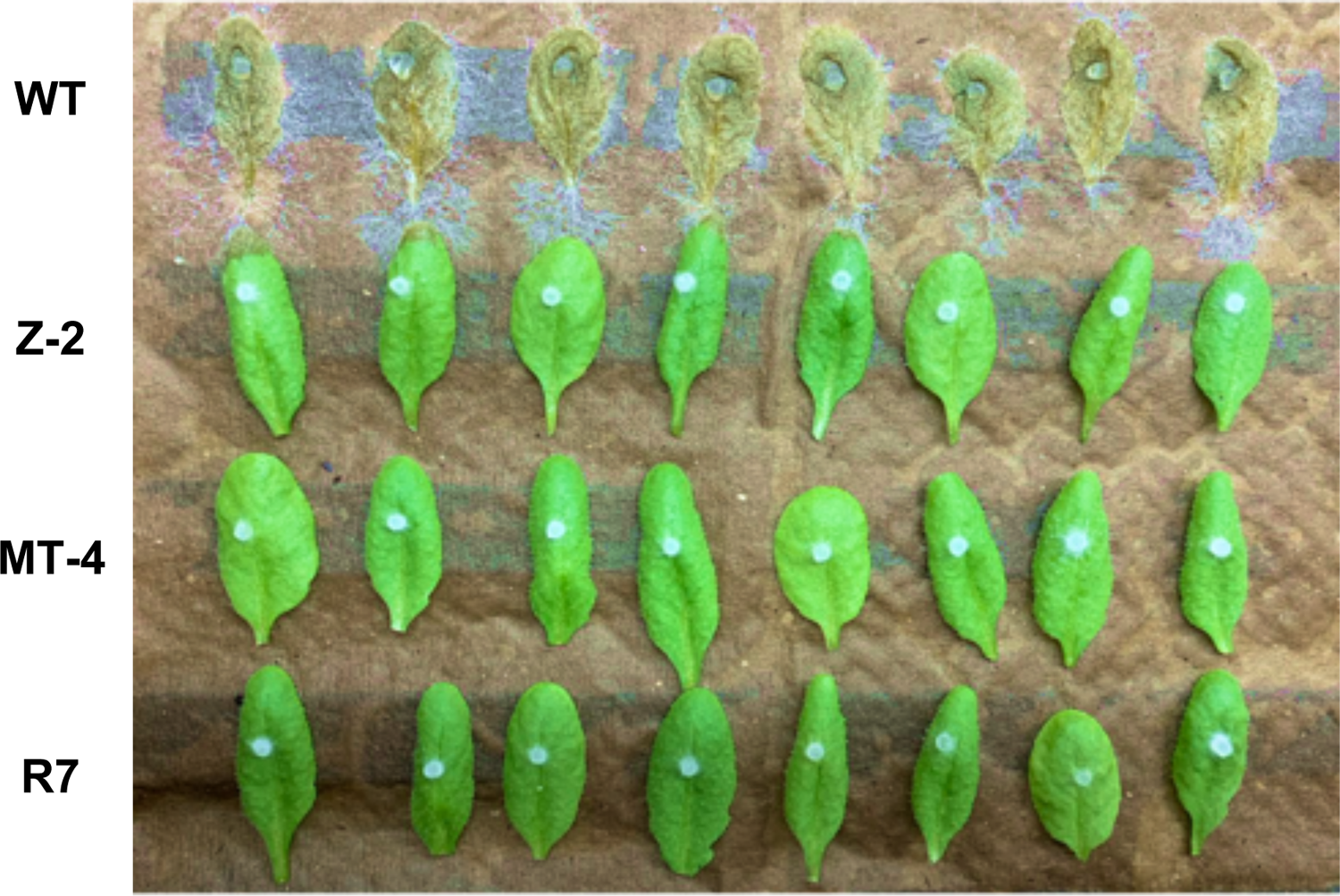
Pathogenicity test of *S. sclerotiorum* mutants Z-2, MT-4, and R7 on *A. thaliana*. Mycelial agar plugs of the indicated *S. sclerotiorum* strains were inoculated onto detached *A. thaliana* Col-0 leaves for pathogenicity test. The picture was taken at 4 dpi.

**Supplemental Figure S2.**
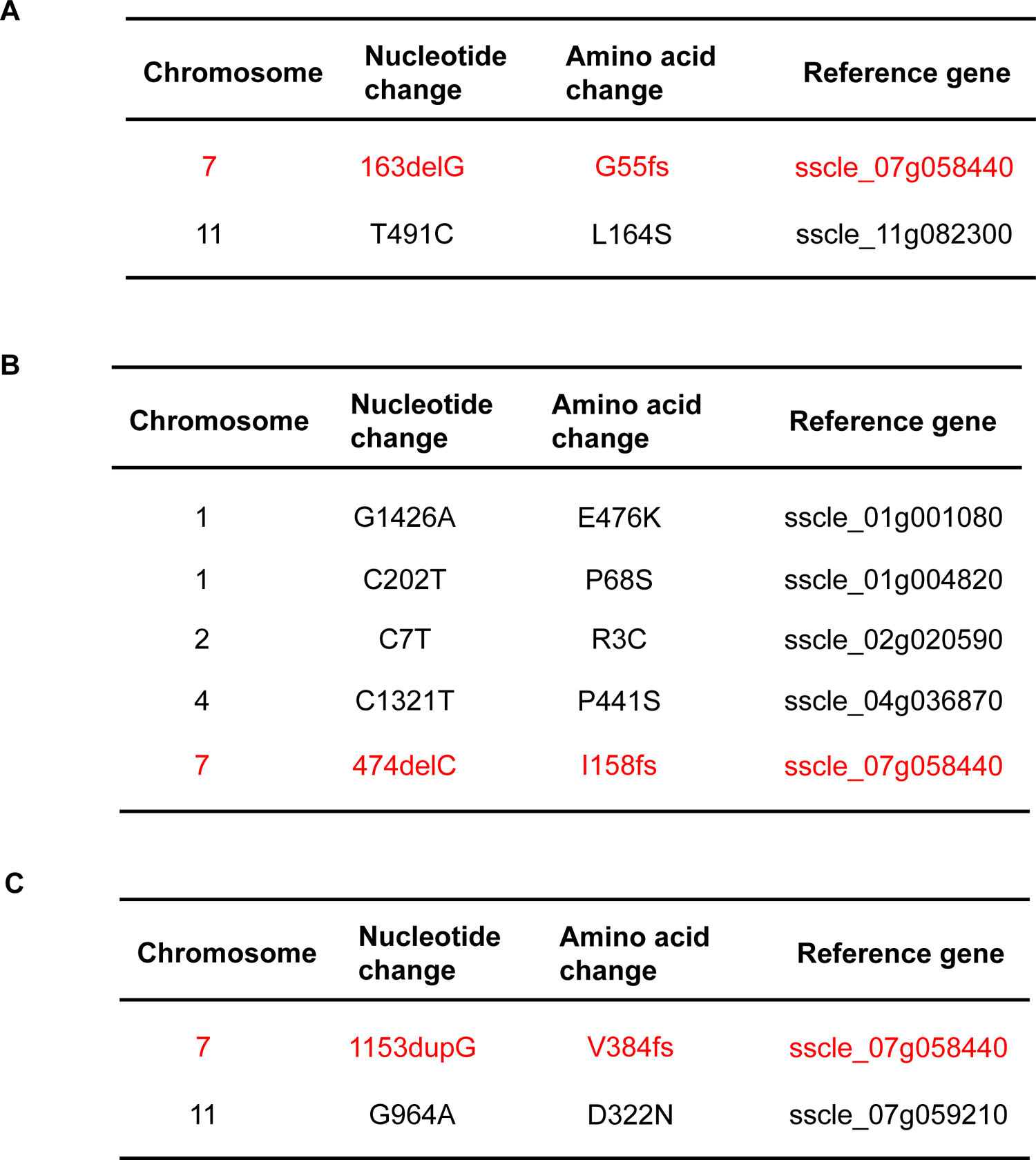
Candidate mutations of Z-2, MT-4, and R7 from NGS data analysis. The most likely gene *sscle_07g058440* from the three mutants is highlighted in red. del: deletion; fs: frameshift; dup: duplication.

**Supplemental Figure S3.**
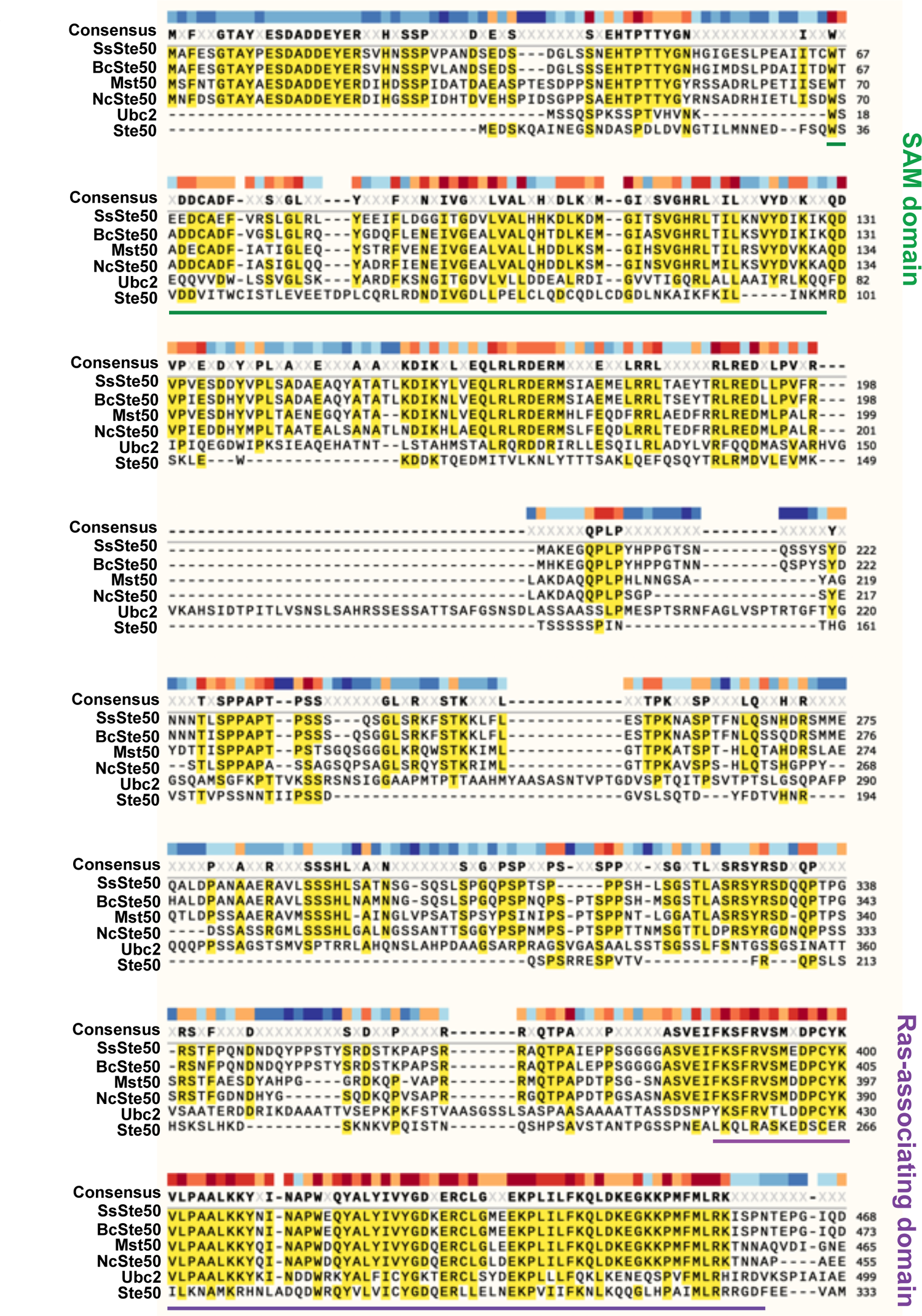
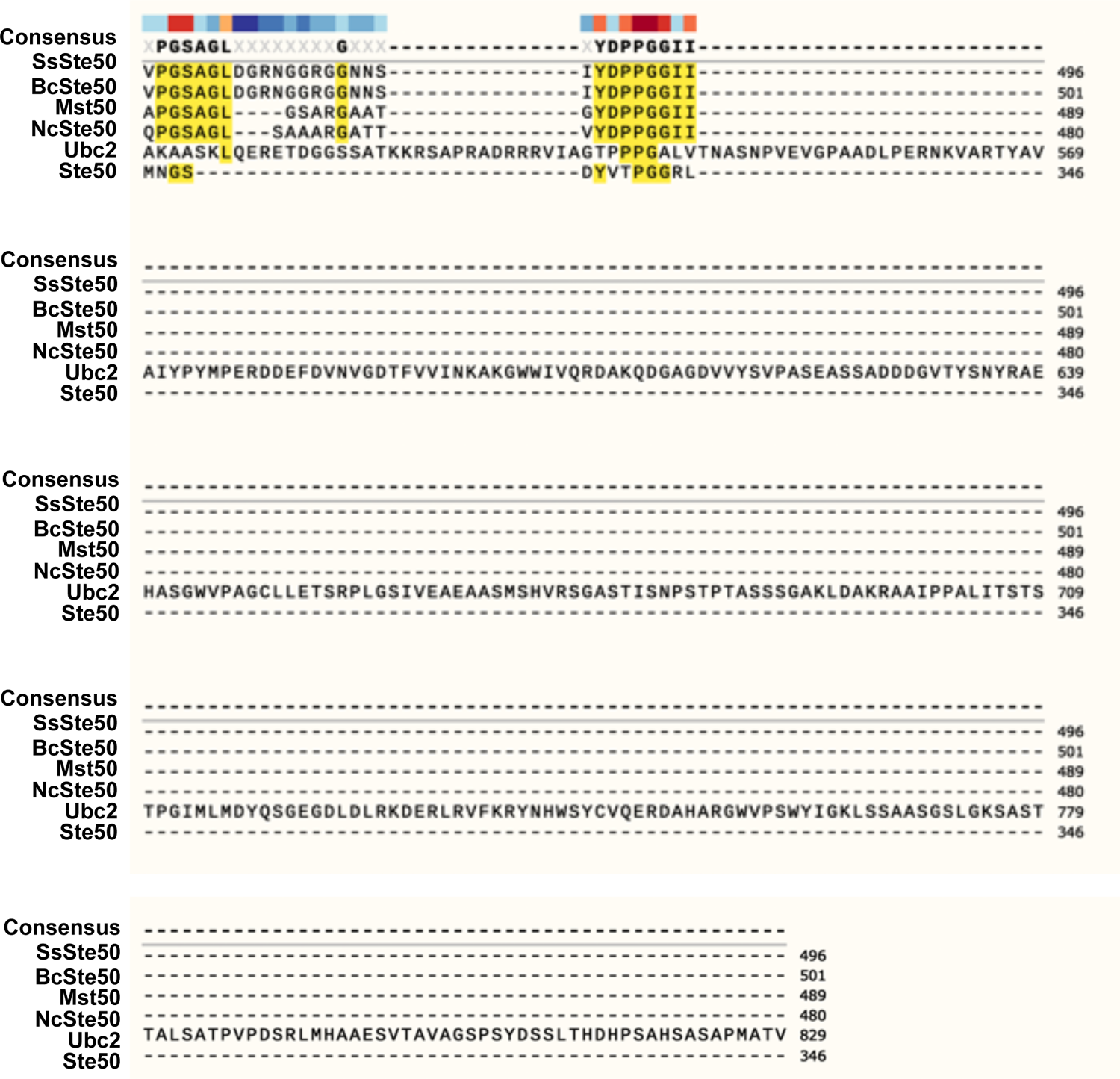
Protein sequence alignment of SsSte50 with its closest homologs in different fungal species. The closest SsSte50 homologs in *Botrytis cinerea* (BcSte50, NCBI ID: XP_001553945.1), *Magnaporthe oryzae* (Mst50, NCBI ID: XP_003712743.1), *Neurospora crassa* (NcSte50, NCBI ID: XP_956774.1), *Ustilago maydis* (Ubc2, NCBI ID: XP_011391983.1), *Saccharomyces cerevisiae* (Ste50, NCBI ID: AJQ34958.1) were used for the protein sequence alignment analysis. SAM domain and Ras-associating domain are underlined in green and purple respectively.

**Supplemental Figure S4.**
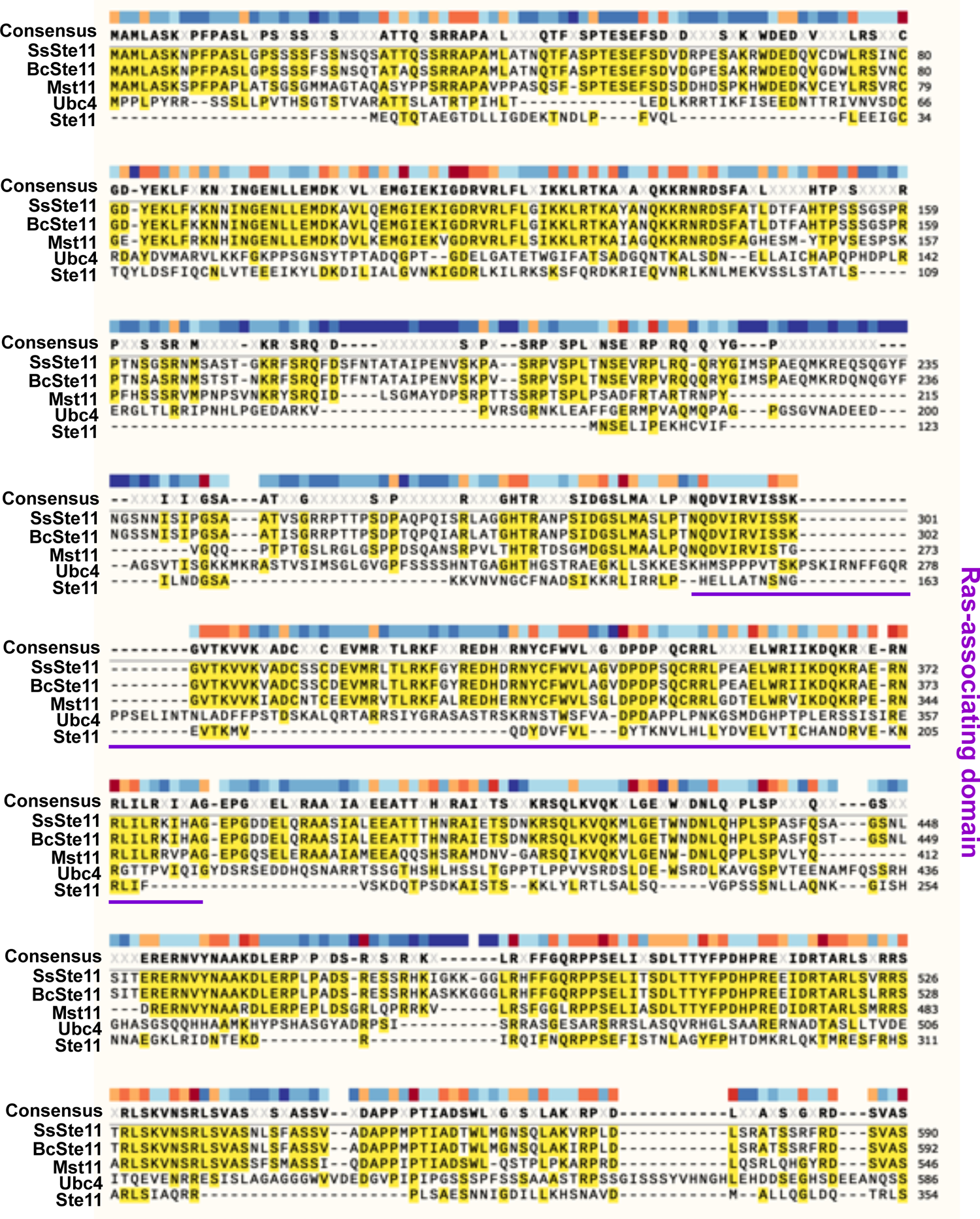
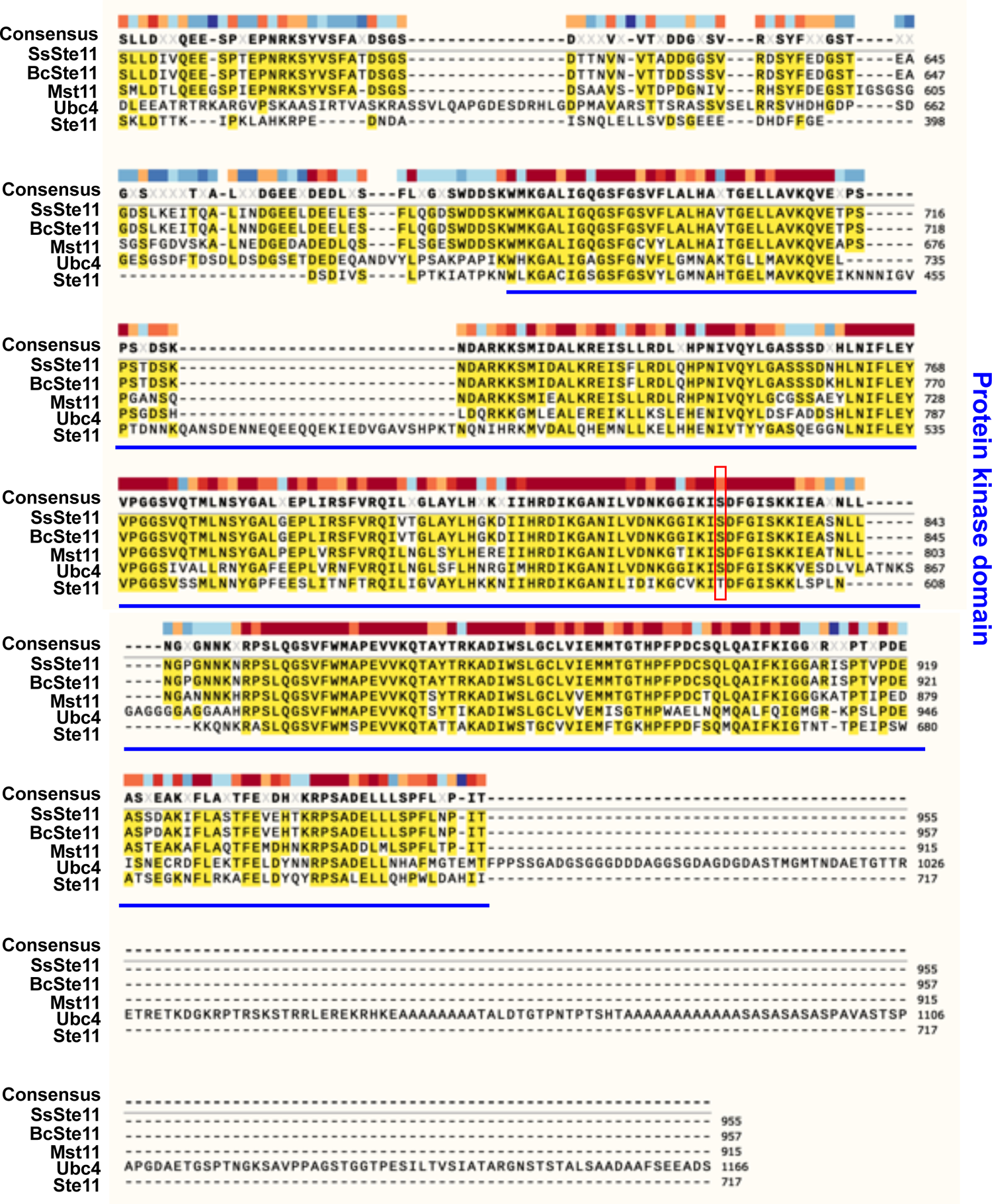
Protein sequence alignment of SsSte11 with its closest homologs in different fungal species. The closest SsSte11 homologs in *B. cinerea* (BcSte11, NCBI ID: XP_024547605.1), *M. oryzae* (Mst11, NCBI ID: XP_003717243.1), *U. maydis* (Ubc4, NCBI ID: AAF86841.1), *S. cerevisiae* (Ste11, NCBI ID: NP_013466.1) were used for the protein sequence alignment analysis. Ras- associating domain is underlined in purple. The conserved protein kinase domain is underlined in blue. The amino acid changed in M30 mutant (*Ssste11-1*) is highlighted in a red box.

**Supplemental Figure S5.**
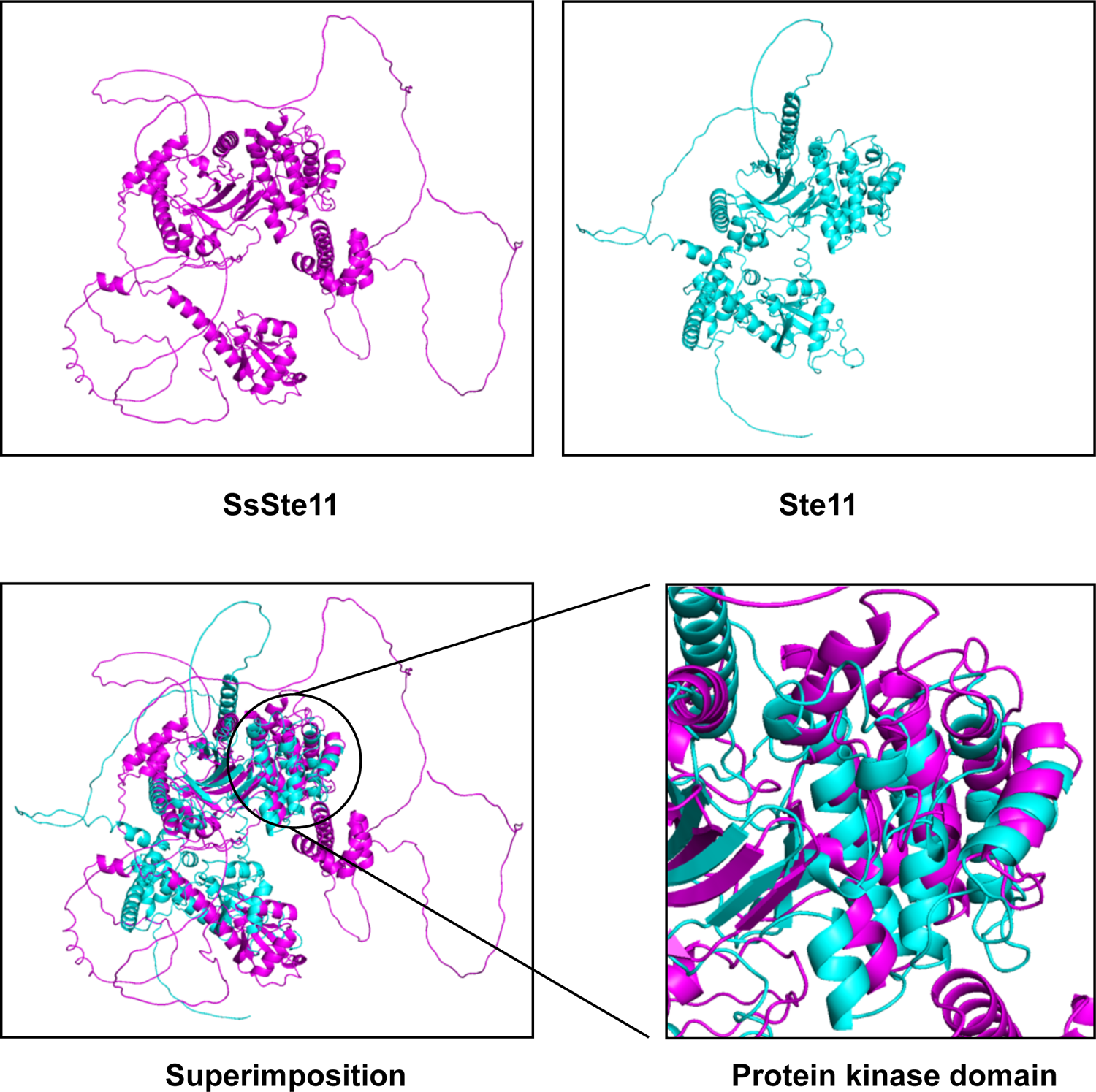
Structural superimposition of SsSte11 and Ste11. AlphaFold-predicted structures of SsSte11 and yeast Ste11 are shown in magenta and cyan respectively. The protein structures were aligned in PyMOL (Schrödinger LLC) using the “super” structural imposition function with an RMSD value of 11.314.

**Supplemental Figure S6.**
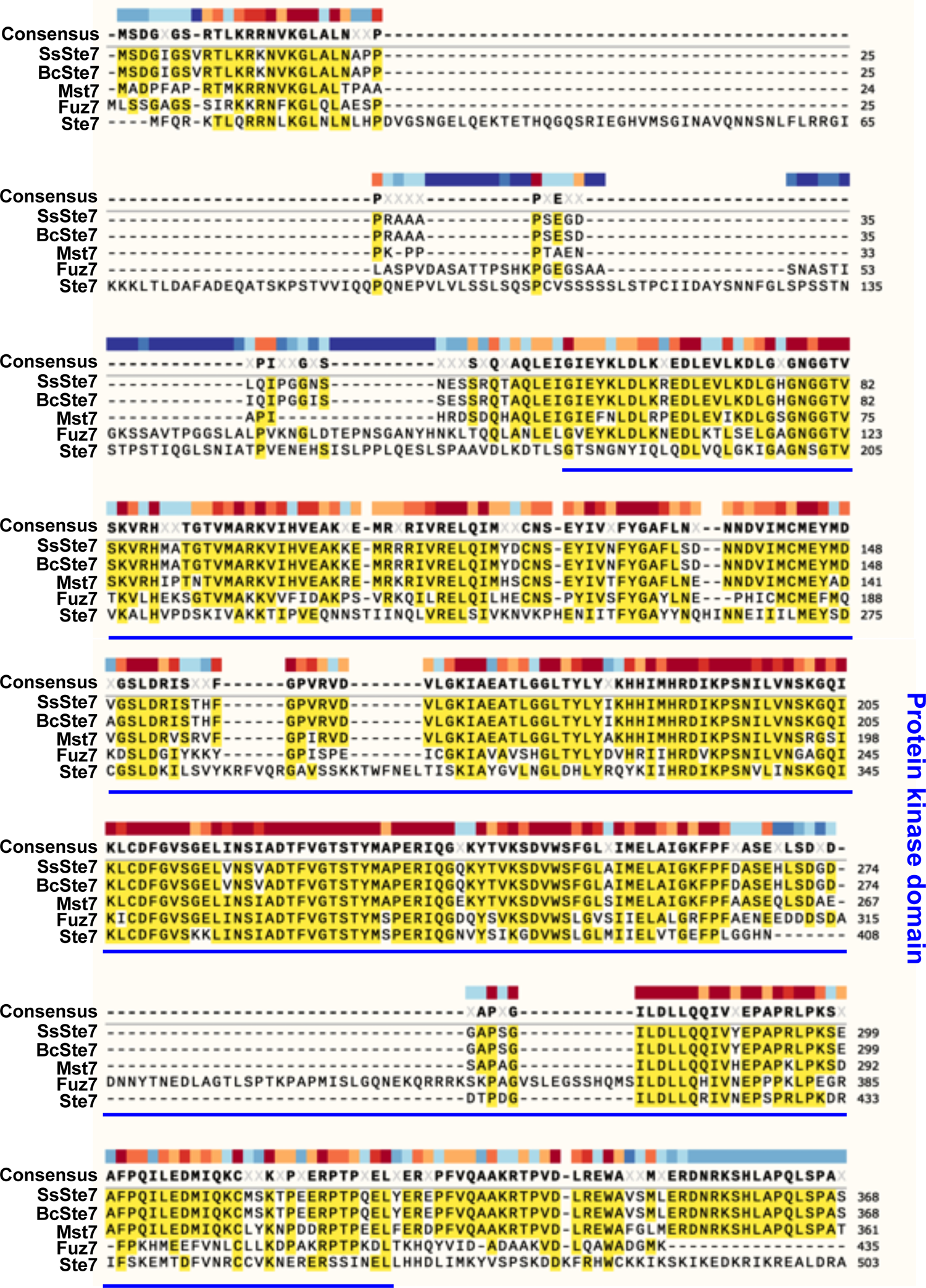
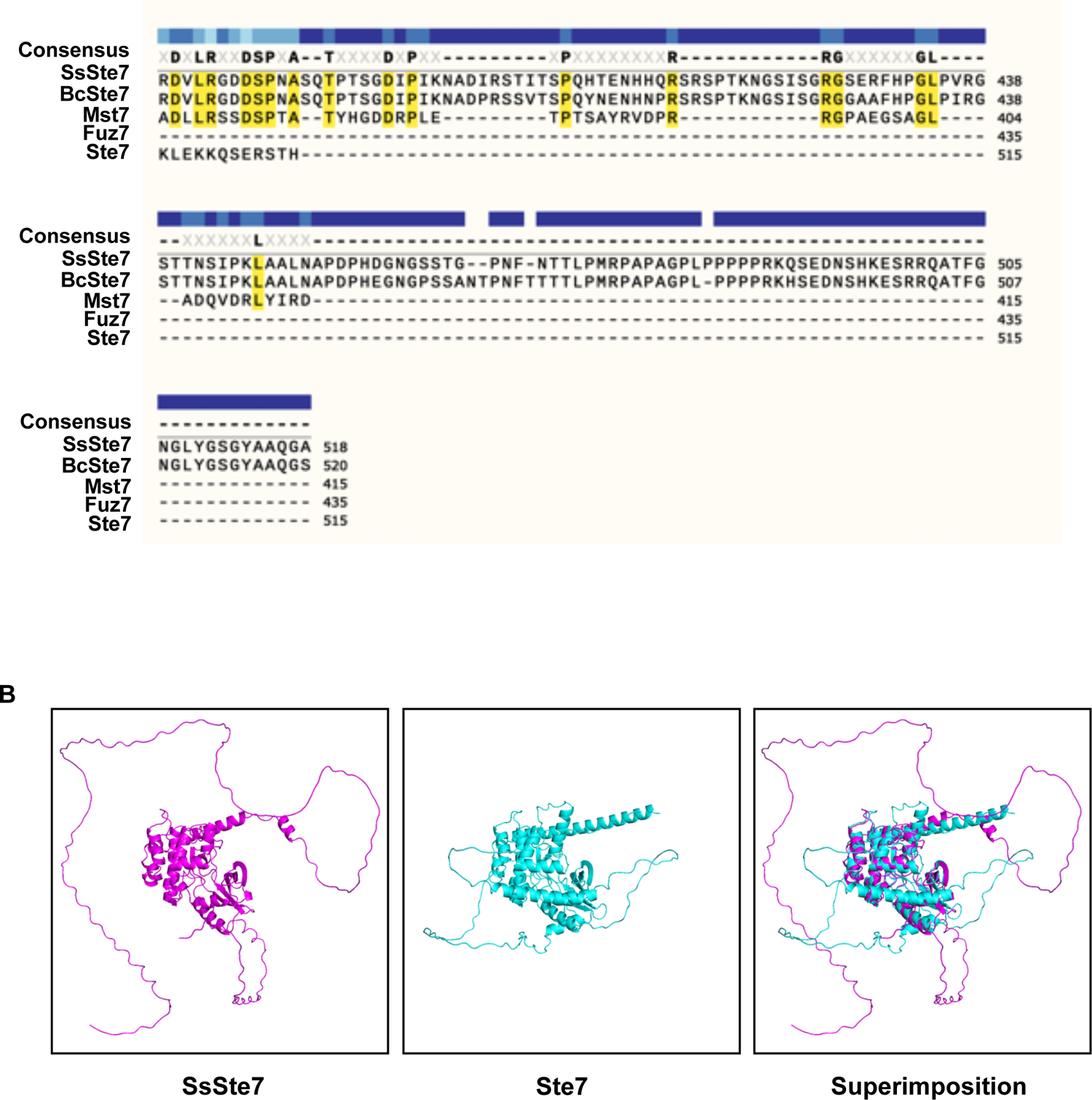
Protein sequence alignment of SsSte11 with its closest homologs in different fungal species. A. The closest SsSte7 homologs in *B. cinerea* (BcSte7, NCBI ID: XP_001557712.1), *M. oryzae* (Mst7, NCBI ID: XP_003718185.1), *U. maydis* (Fuz7, NCBI ID: XP_011387552.1), *S. cerevisiae* (Ste7, NCBI ID: AJP37595.1) were used for the protein sequence alignment analysis. The conserved protein kinase domain is underlined in blue. B. Structural superimposition of SsSte7 and Ste7. AlphaFold-predicted structures of SsSte7 and yeast Ste7 are shown in magenta and cyan respectively. The protein structures were aligned in PyMOL (Schrödinger LLC) using the “super” structural imposition function with an RMSD value of 1.361.

**Supplemental Figure S7.**
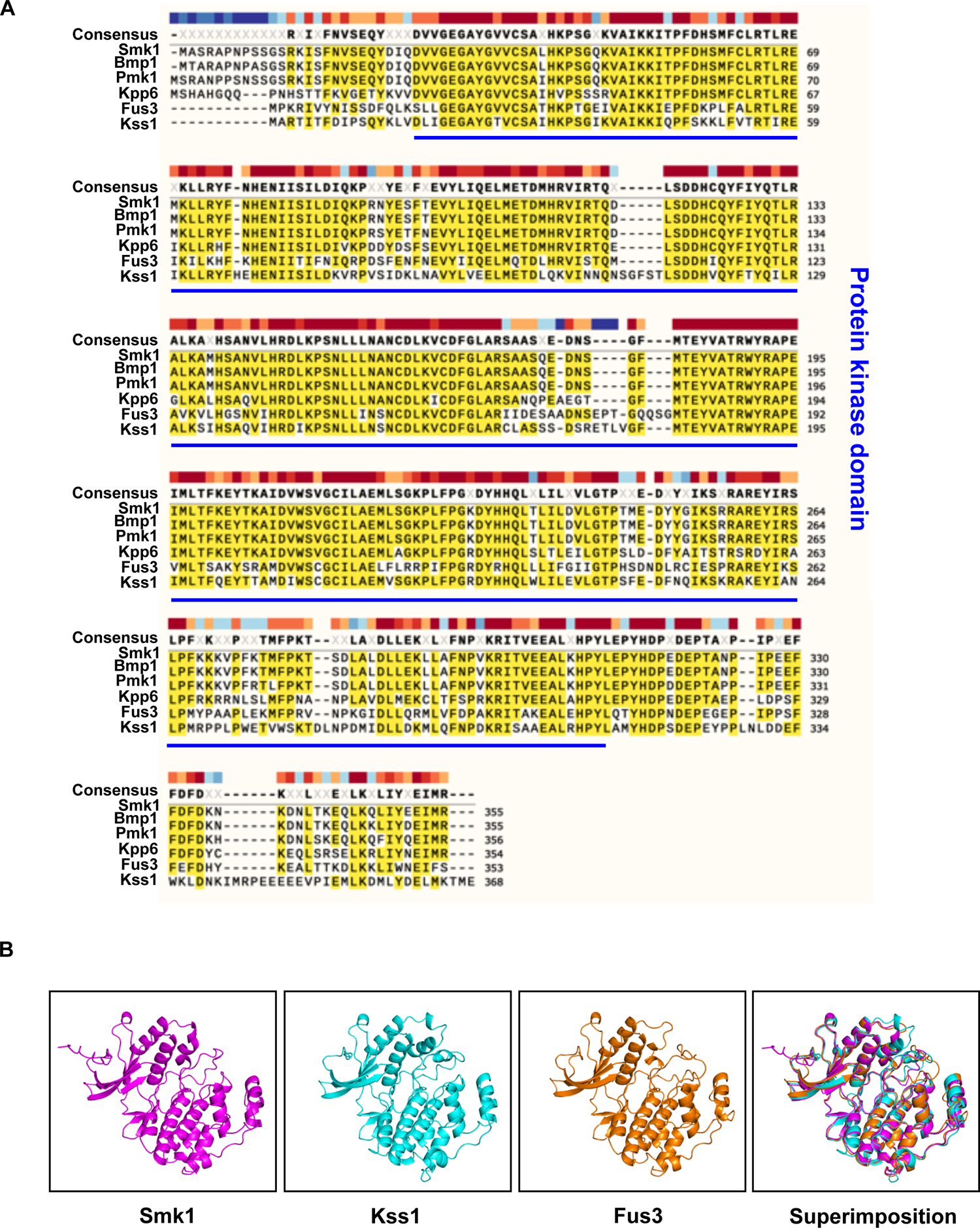
Protein sequence alignment of Smk1 with its closest homologs in different fungal species. A. The closest Smk1 homologs in *B. cinerea* (Bmp1, NCBI ID: XP_024547333.1), *M. oryzae* (Pmk1, NCBI ID: XP_003712175.1), *U. maydis* (Kpp6, NCBI ID: XP_011389711.1), *S. cerevisiae* (Fus3, NCBI ID: AAA34613.1; Kss1, NCBI ID: NP_011554.3) were used for the protein sequence alignment analysis. The conserved protein kinase domain is underlined in blue. B. Structural superimposition of Smk1, Kss1 and Fus3. AlphaFold-predicted structures of Smk1 and yeast Kss1 and Fus3 are shown in magenta, cyan and orange respectively. The protein structures were aligned in PyMOL (Schrödinger LLC) using the “super” structural imposition function with RMSD values of 0.659 (Smk1 with Kss1) and 0.630 (Smk1 with Fus3).

**Supplemental Table S1.**
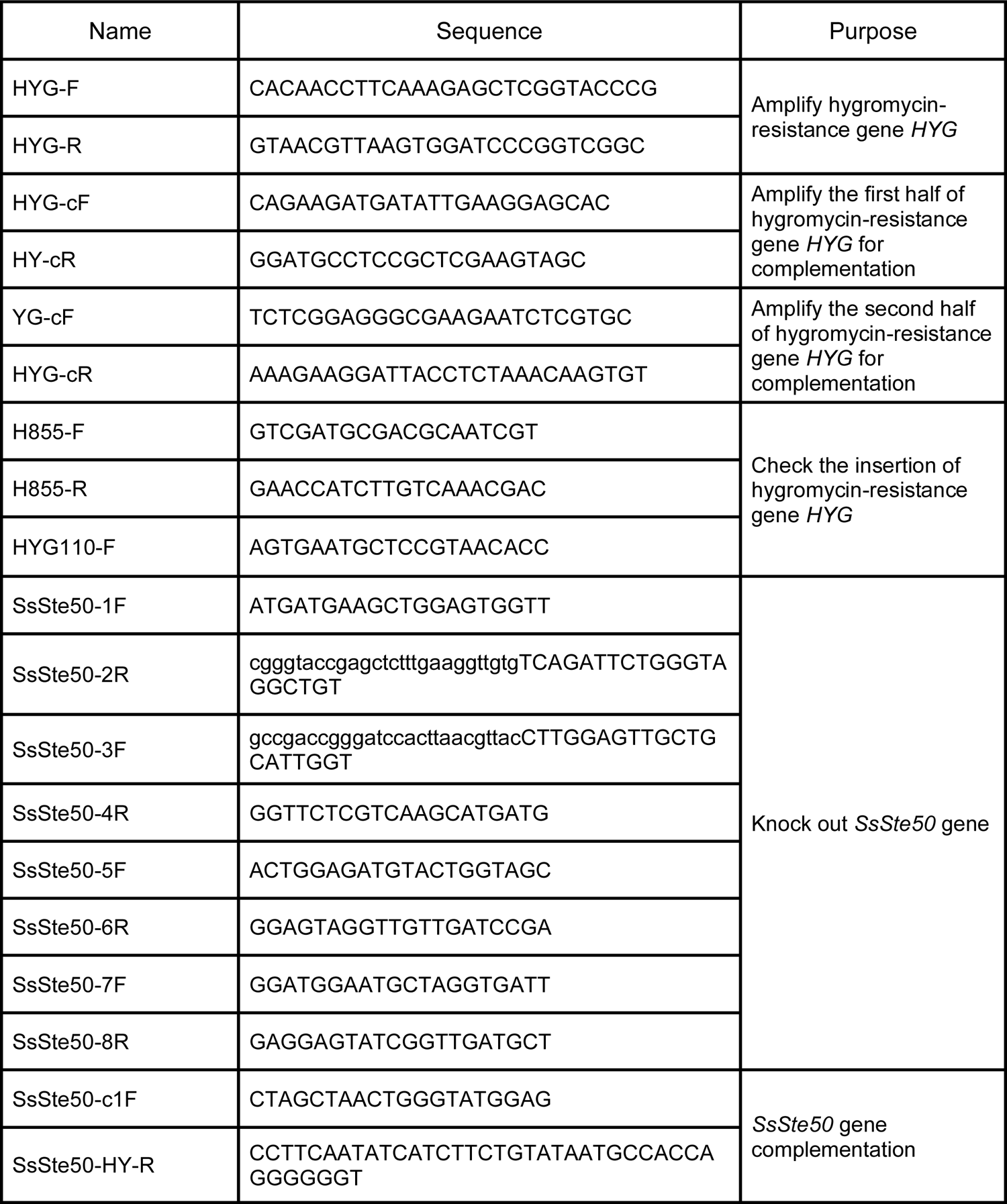
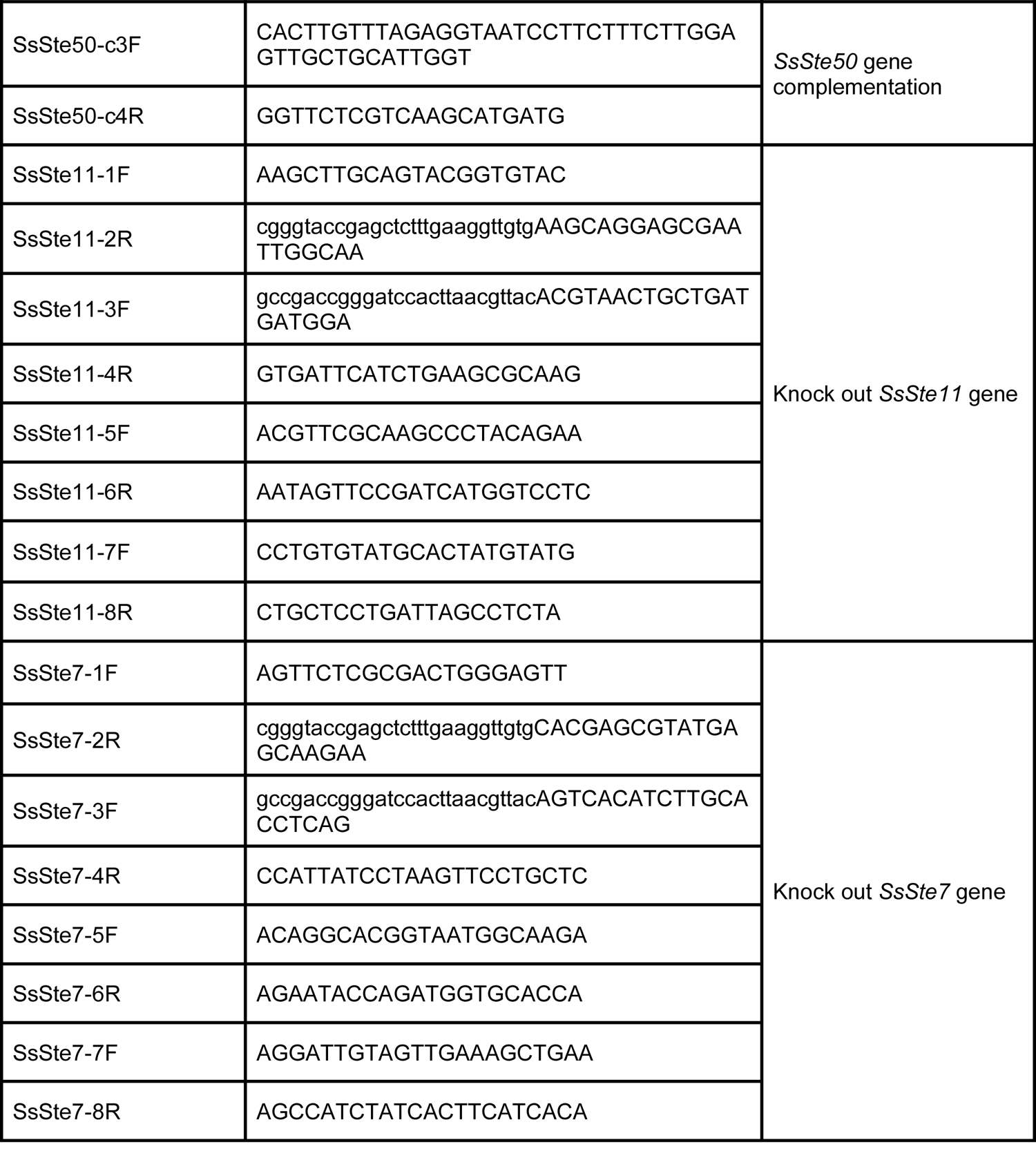
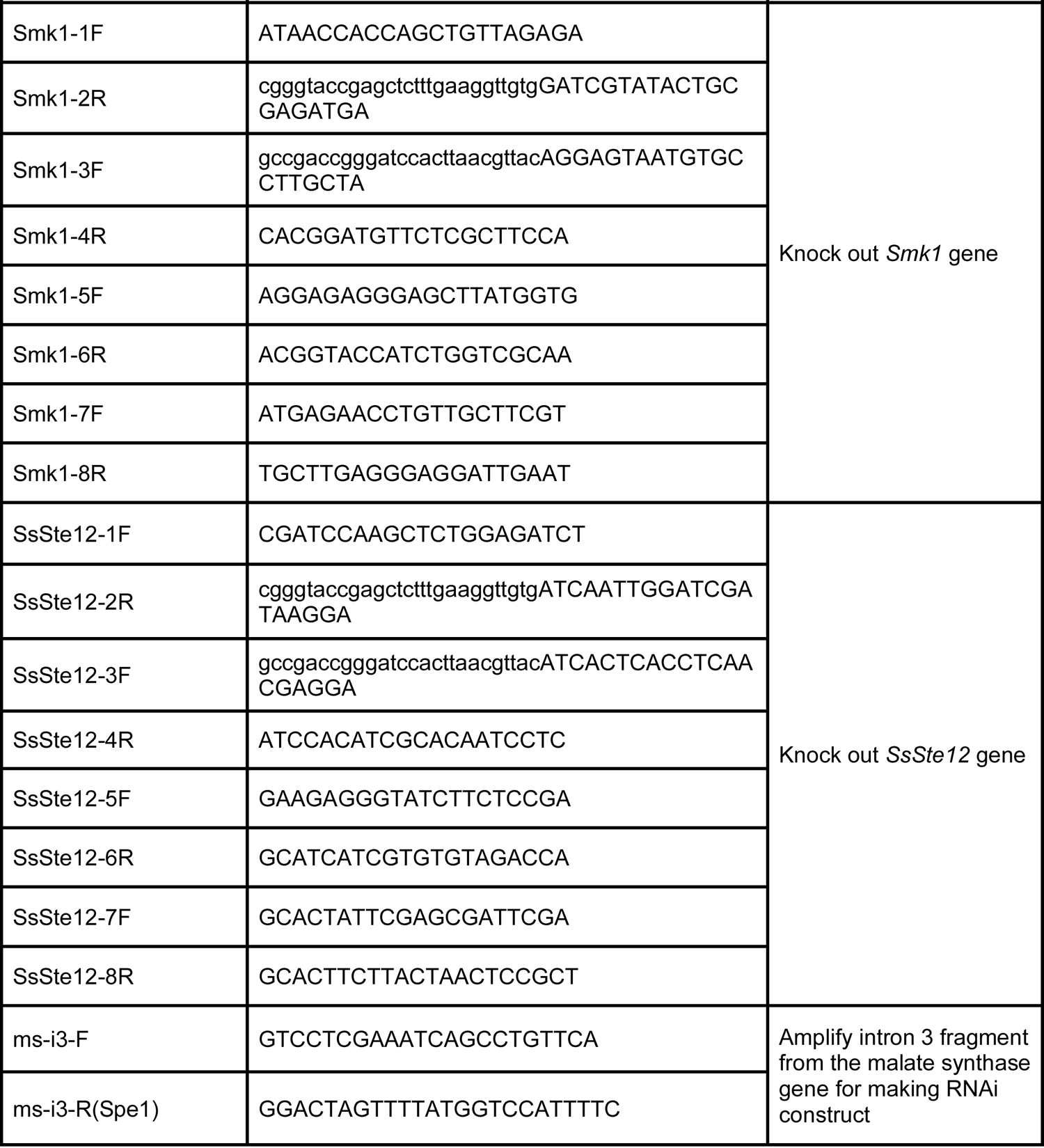
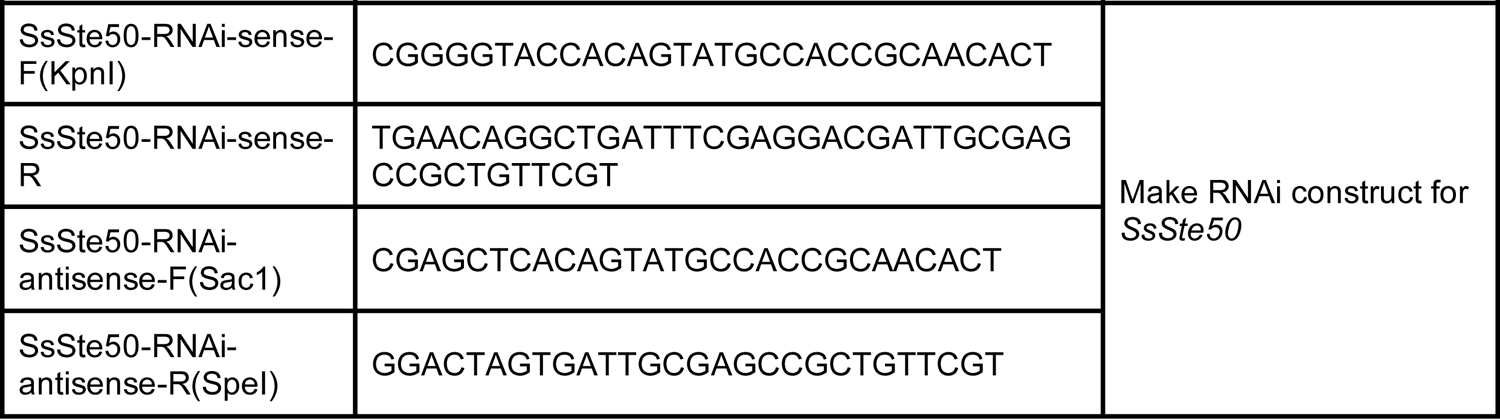
The list of primers used in this study.

## Acknowledgements

We cordially thank Dr. Jeffrey Rollins from University of Florida for generous sharing of *S. sclerotiorum* strain 1980 and the *S. sclerotiorum oah1* mutant; Dr. Daohong Jiang from Huazhong Agricultural University for generous sharing of *pCH-EF-1* vector. This work was supported by funds to X.L. from the Natural Sciences and Engineering Research Council (NSERC) Discovery program of Canada, NSERC-CREATE-PRoTECT, and the Canadian Foundation for Innovation (CFI). L.T. and Y.X. are partly supported by China Scholarship Council (CSC) scholarships.

## Author Contributions

L.T. contributed to conceptualization, data curation, validation, investigation, methodology, writing - original draft, project administration; J.L. contributed to validation, methodology and writing - original draft; Y.X. identified the M30 mutant from the forward genetic screen and contributed to methodology; Y.Q. identified the MT-4 mutant. X.L. contributed to conceptualization, data curation, formal analysis, supervision, funding acquisition, project administration, writing - revisions. All authors reviewed the manuscript.

## Competing financial interests

The authors declare no competing financial interests.

**Correspondence and requests for materials** should be addressed to XL.

